# Transcriptional and Cellular Response of hiPSC-derived Microglia-Neural Progenitor Co-Cultures Exposed to IL-6

**DOI:** 10.1101/2023.12.21.572748

**Authors:** Amalie C. M. Couch, Amelia M. Brown, Catarina Raimundo, Shiden Solomon, Morgan Taylor, Laura Sichlinger, Rugile Matuleviciute, Deepak P. Srivastava, Anthony C. Vernon

**Affiliations:** Department of Basic and Clinical Neuroscience, Institute of Psychiatry, Psychology and Neuroscience, King’s College London, London, UK; MRC Centre for Neurodevelopmental Disorders, King’s College London, London, UK; Faculty of Life Sciences and Medicine, King’s College London, London, UK

**Keywords:** IL-6, neurodevelopment, human induced-pluripotent stem cells, microglia, neural progenitor cells

## Abstract

Elevated interleukin (IL-)6 levels during prenatal development have been linked to increased risk for neurodevelopmental disorders (NDD) in the offspring, but the mechanism remains unclear. Human-induced pluripotent stem cell (hiPSC) models offer a valuable tool to study the effects of IL-6 on features relevant for human neurodevelopment *in vitro*. We previously reported that hiPSC-derived microglia-like cells (MGLs) respond to IL-6, but neural progenitor cells (NPCs) in monoculture do not. Therefore, we investigated whether co-culturing hiPSC-derived MGLs with NPCs would trigger a cellular response to IL-6 stimulation via secreted factors from the MGLs. Using N=4 donor lines without psychiatric diagnosis, we first confirmed that NPCs can respond to IL-6 through trans-signalling when recombinant IL-6Ra is present, and that this response is dose-dependent. MGLs secreted soluble IL-6R, but at lower levels than found *in vivo* and below that needed to activate trans-signalling in NPCs. Whilst transcriptomic and secretome analysis confirmed that MGLs undergo substantial transcriptomic changes after IL-6 exposure and subsequently secrete a cytokine milieu, NPCs in co-culture with MGLs exhibited a minimal transcriptional response. Furthermore, there were no significant cell fate-acquisition changes when differentiated into post-mitotic cultures, nor alterations in synaptic densities in mature neurons. These findings highlight the need to investigate if trans-IL-6 signalling to NPCs is a relevant disease mechanism linking prenatal IL-6 exposure to increased risk for psychiatric disorders. Moreover, our findings underscore the importance of establishing more complex *in vitro* human models with diverse cell types, which may show cell-specific responses to microglia-released cytokines to fully understand how IL-6 exposure may influence human neurodevelopment.

## Introduction

Human birth cohort studies have linked elevated maternal interleukin (IL)-6 concentrations across gestation with aberrant structural and functional brain connectivity in the offspring, as measured by MRI, which in turn are associated with abnormal cognitive development and increased risk for neurodevelopmental disorders (NDDs) such as autism spectrum condition (ASC) and schizophrenia (SZ) (Graham *et al*., 2018; Rudolph *et al*., 2018; Rasmussen *et al*., 2019; Allswede *et al*., 2020). Although these cohort studies demonstrate a correlation between elevated prenatal IL-6 and NDD incidence, additional studies at a molecular and cellular level are required to elucidate the molecular mechanisms by which this risk is conferred. Much of the cellular level research that links prenatal IL-6 insults with adverse neurodevelopmental effects in the offspring has, however, been conducted in rodent models (Samuelsson *et al*., 2006; Smith *et al*., 2007; Ozaki *et al*., 2020; Mirabella *et al*., 2021), which may have different brain development trajectories (Eze et al., 2021) For example, transient elevation of IL-6 during a critical period of neurodevelopment can have enduring effects on the formation of glutamatergic synapses in the mouse hippocampus with consequences for mouse brain connectivity as measured by resting state fMRI (Samuelsson *et al*., 2006; Mirabella *et al*., 2021). Nonetheless, investigating the influence of IL-6 on early neurodevelopmental processes using human-led models is crucial to understand its role in increasing the risk of NDDs in offspring (Bayés *et al*., 2011; Südhof, 2017; Kuljis *et al*., 2019).

As previously reviewed, IL-6 signalling involves both the IL-6 receptor (IL-6Ra) and IL-6 signal transducer (IL-6ST) subunits, with IL-6 binding to IL-6Ra triggering a conformational change that activates the JAK/STAT pathway through either cis-signalling (cell surface IL-6Ra expression) or trans-signalling (via secreted soluble IL-6Ra, sIL-6Ra) (Rose-John, 2001, 2012, 2018; Wolf, Rose-John and Garbers, 2014). Trans-signalling via sIL-6Ra enables activation of cells lacking the full receptor and may be implicated in pro-inflammatory effects on non-glial cells, contrasting cis-signalling’s proposed anti-inflammatory effects (Rose-John, 2001, 2012; Rose-John *et al*., 2009; Scheller *et al*., 2011; Campbell *et al*., 2014; Wolf, Rose-John and Garbers, 2014). This mechanism is known to occur in the CNS, demonstrated when blocking trans-signalling with soluble IL-6ST in the CNS of adult rodents, which resulted in reduced astrogliosis and microgliosis, blood-brain barrier leakage, vascular proliferation, neurodegeneration, as well as impaired neurogenesis, caused by overexpression of IL-6 (Campbell *et al*., 2014). However, the consequences of trans-IL-6 signalling during human development remains unclear.

Microglia are central to the developing brain’s response to IL-6. In mid-gestation (GW17-18) human neocortex tissue (Polioudakis *et al*., 2019), as well as in the developing human telencephalon (Florio *et al*., 2015; Zhang *et al*., 2016; Nowakowski *et al*., 2017), microglia are the leading cell type expressing IL-6Ra, with expression levels being higher than in excitatory neurons, intermediate progenitors, interneurons, radial glia, mitotic progenitors, oligodendrocyte precursors, endothelial cells and pericytes. Importantly, initiation of either IL-6 trans- or cis-signalling may be dependent on the expression ratios of the receptor complex’s subunits (Reeh *et al*., 2019). However, the direct link between sIL-6Ra secretion by microglia and enhanced trans-signalling in non-glial cells (Ferreira *et al*., 2013; Campbell *et al*., 2014), as well as its implications in a neurodevelopment system remain uncertain, especially in a human-relevant model. This knowledge gap underscores the value of using human-induced pluripotent stem cells (hiPSCs) to model human foetal brain development (Eze *et al*., 2021). In agreement with primary tissue studies (Florio *et al*., 2015; Zhang *et al*., 2016; Nowakowski *et al*., 2017; Polioudakis *et al*., 2019), hiPSC-derived NPCs express IL-6ST, yet they lack IL-6Ra expression (Couch *et al*., 2023; Sarieva, Hildebrand, *et al*., 2023). This renders them unresponsive to IL-6 in mono-culture without the addition of sIL-6Ra necessary for trans-signalling (Couch *et al*., 2023; Sarieva, Hildebrand, *et al*., 2023). To circumvent this limitation, previous *in vitro* studies by using hyper-IL-6, a synthetic construct combining sIL-6Ra and IL-6 linked by a flexible peptide, enabling NPCs to respond to IL-6 through trans-signalling (Sarieva, Hildebrand, *et al*., 2023). Recent investigations using a 3D human forebrain organoid, which lacked microglia, demonstrated that exposure to hyper-IL-6 disproportionately affected radial glia cells, leading to an increased presence of this cell type in the organoid (Sarieva, Kagermeier, *et al*., 2023). Notably, chronic 5-day hyper-IL-6 exposure also resulted in an increase in upper-layer excitatory neurons and abnormalities in neuronal migration and positioning, suggesting that forced IL-6 trans-signalling can disrupt the standard process of cortical development and organisation (Sarieva, Kagermeier, *et al*., 2023). These data provide human-relevant evidence that IL-6 signalling in developing forebrain organoids can change typical cellular and molecular phenotypes.

Although hyper-IL-6 effectively demonstrates that NPCs are capable of IL-6 trans-signalling, it does not replicate the hypothesised physiological trans-signalling process found *in vivo.* Hyper-IL-6 forces cells to respond via trans-signalling under any condition, despite the fact there could be an upstream signalling regulation mechanism governed by non-NPC cell types. Our previous research indicates that hiPSC-derived microglia-like cells (MGLs) can secrete sIL-6Ra, addressing the absence of this subunit in hiPSC-derived NPCs (Couch *et al*., 2023). Consequently, a co-culture system of MGLs and developing cortical NPCs may offer a more physiologically relevant model to study the acute IL-6 responses of both cell types, providing a closer approximation to physiological conditions than hyper-IL-6. In this study, we employ hiPSC-derived MGLs and NPCs from the same donors for co-culture, ensuring a consistent genetic background within each culture, and expose them to acute IL-6 stimulation for 24 hours. Employing a trans-well co-culture system allows for the analysis of soluble factors, minimises potential confounding impact from the experimental handling of microglia (Cadiz *et al*., 2022), and precludes the need for more invasive separation methods like fluorescence-activated cell sorting (FACS) (Park *et al*., 2020). The overall study aim was to ascertain the cellular and molecular changes of both NPCs and MGLs post-IL-6 exposure. We hypothesised that: (1) co-culturing with MGLs will facilitate acute NPC response to IL-6 via trans-signalling mediated by sIL-6Ra, or additional cytokines secreted by MGLs in response to IL-6; (2) this interaction will modify the transcriptomes of both MGLs and NPCs, as identified through bulk RNAseq, revealing key molecular pathways; (3) early exposure of NPCs to cytokines will influence their cell fate determination and synaptogenesis upon differentiation into post-mitotic cortical neurons.

## Methods

### Cell Culture

#### hiPSCs

For the derivation of human induced pluripotent stem cells (hiPSCs), participants were recruited, and methods carried out in accordance with the ‘Patient iPSCs for Neurodevelopmental Disorders (PiNDs) study’ (REC No 13/LO/1218). Informed consent was obtained from all subjects for participation in the PiNDs study. Ethical approval for the PiNDs study was provided by the NHS Research Ethics Committee at the South London and Maudsley (SLaM) NHS R&D Office. Human iPSCs were generated from a total of four lines donated by three males and one female with no history of neurodevelopmental or psychiatric disorders (Supplementary Table 1). All hiPSC donor lines used in this study have been previously characterised, as described elsewhere (Warre-Cornish *et al*., 2020; Adhya *et al*., 2021; Pantazis *et al*., 2022; Couch *et al*., 2023). As such and following the International Society for Stem Cell Research (ISSCR) standards for previously published lines, we performed karyotyping and confirmed the expression of pluripotency markers (Supplementary Figure 1). The cells were grown in hypoxic conditions on Geltrex™ (Life Technologies; A1413302) coated 6-well NUNC^TM^ plates in StemFlex medium (Gibco, A3349401) exchanged every 48 hours. For passaging, cells were washed with HBSS (Invitrogen; 14170146) and then passaged by incubation with Versene (Lonza; BE17-711E), then plated in fresh StemFlex onto fresh Geltrex-coated 6-well NUNC^TM^ plates.

#### Microglia-like and Neural Progenitor Cell Co-culture

Microglia-like cells (MGL) differentiation from hiPSCs was performed following a previously published protocol (van Wilgenburg *et al*., 2013; Haenseler *et al*., 2017), and successfully replicated by our group (Couch *et al*., 2023). Simultaneously but in separate cultures, hiPSCs from the same lines were neuralised towards neural progenitor cells (NPCs) using a modified dual SMAD inhibition protocol to develop a mixture of excitatory and inhibitory forebrain neuronal subtypes (Shi, Kirwan and Livesey, 2012; Warre-Cornish *et al*., 2020; Adhya *et al*., 2021; Bhat *et al*., 2022; Couch *et al*., 2023), and were characterised for Nestin and β-tubulin III (TUBB3, antibody name TUJ1) expression using immunocytochemistry (Supplementary Figure 2). MGL progenitors were seeded directly from the embryoid body factory onto 0.4 µm Transparent PET Membrane Permeable Support Cell Culture Inserts (Falcon®; 353090) and allowed to differentiate for 12 days, after which they were assembled on top of the NPCs that had been differentiating in parallel for 16 days (Supplementary Figure 3). This permitted the NPCs to settle after neural passaging the day before on day 15 of their differentiation. Both cell types were then cultured together for 48h in co-culture media (1X N2 supplement, 2mM Glutamax, 100ng/ml IL-34 and 10ng/ml GM-CSF), to allow the two cell types to “equilibrate”. Specifically, during their initial optimization trials for establishing the hiPSC to MGL differentiation method, Haenseler *et al*. (2017) raised concerns regarding the presence of corticosterone, superoxide dismutase, and catalase compounds in the B27 supplement. Therefore, B27 was removed from co-culture medium to guarantee compatibility with iPSC-derived cortical neurons while preserving microglial function (Haenseler *et al*., 2017). The co-cultures were then exposed to 100ng/ml IL-6 or vehicle control (sterile water with 100 pM acetic acid) in fresh media, for 24h (Supplementary Figure 3). This meant stimulation on the microglial differentiation day 14, and neural progenitor differentiation day 18 (Couch *et al*., 2023). The differentiation timepoints and 100ng/ml IL-6 dosage were based on previously published data that examined the IL-6 dose-response in day 14 MGLs (Couch *et al*., 2023). After 24h of IL-6 or vehicle exposure, MGLs and NPCs were reseparated. All MGLs were collected for RNAseq, half NPCs were collected for RNAseq, and half were terminally differentiated into post-mitotic neurons as follows (Supplementary Figure 3).

#### Post-mitotic Cortical Neuron Differentiation

After 24h of IL-6 stimulation in co-culture with MGLs, half the NPCs were subsequently terminally differentiated into post-mitotic cortical neurons. NPCs to be terminally differentiated were passaged three days after IL-6 stimulation (D21) into 50ng/ml poly-l-ornithine (Gibco; A3890401) and 20ng/ml laminin (Sigma; L2020) coated 96-well Perkin Elmer CellCarrier Ultra plates, at a density of 30,000 cells/well in 1X B27 + 10µM DAPT + 10µM Rock inhibitor and incubated (37°C; 5% CO_2_; 20% O_2_). After 24 hours, medium was replaced with Rock inhibitor-free medium every 24 hours until D28. After this time frame, the notch inhibitor DAPT was removed from the medium and cells were fed with half 1X B27 medium exchanges every 7 days.

### Immunoblotting

Protein samples were collected by scraping in ice cold RIPA Lysis and Extraction Buffer (Thermo Scientific; 89900), with Phosphatase Inhibitor Cocktail 3 (1:100 Sigma; P0044), AEBSF (1 mM), Leupeptin (10 µg/ml) and Aprotinin (10 µg/ml). Cells were then lysed by sonication at 40% for 10 pulses, then centrifuged to pellet out insoluble materials at 20,000 x *g* for 15 min at 4 °C, with proteins collected in the supernatant. Protein concentration was quantified using the Pierce™ BCA protein assay kit (Thermo-Fisher; 23227). In preparation for SDS-PAGE separation, protein samples were denatured in 2x Laemmli Sample Buffer (BioRad; 1610737EDU) with 5% 2-Mercaptoethanol and boiled at 95°C for 5min. 2 µg of each protein sample was loaded into 10% acrylamide gels, alongside 5 µl of the Dual Color (BioRad;1610374) standard marker. Gels were run at 20mA for approximately 20 min, then increased to 100V until the samples reached the bottom of the unit (∼90min). Separated samples were transferred to a PVDF membrane and run overnight at 78mA at 4°C. Blots were blocked in 5% BSA TBS-T for 1 hour at RT with agitation. Antibodies (pSTAT3-Y705 Cell Signalling #9138S, total-STAT3 Cell Signalling #30835S, Phospho-p44/42 MAPK (pERK1/2-T202/Y204) Cell Signalling #9101, p44/p42 MAPK (total ERK1/2) Cell Signalling #4696S, GAPDH abcam #ab9484 and Beta-Actin Invitrogen #MA5-15739) were diluted in blocking buffer; primary antibody incubation occurred overnight at 4°C with agitation, and secondary antibody incubation at RT for 1 hour with agitation. Washes between antibody probes occurred in TBS-T at three 15min intervals. For visualization, ECL Western Blotting Substrate (GE Healthcare; RPN2106) was incubated on the blot at RT for 5min before image capture by the Bio-Rad Molecular Imager^®^ Gel Doc™ XR System. Blots were probed with antibodies directed towards the phosphorylated protein first, then total protein and finally the loading control beta-actin or GAPDH. Blots were stripped between each staining session by incubating the membrane in Restore^TM^ Stripping Buffer (Thermo Fischer Scientific; 21059) for 20mins at RT, then rinsing with TBS-T three times before moving onto re-blocking with 5% BSA TBS-T to commence the next probing phase. Blot signals were quantified using ImageStudioLite (LI-COR, version 5.2.5), by defining an identical rectangular region of interest around each signal band and measuring the median signal value. Background correction was then performed using the automatic background detection toolkit provided with the aforementioned software. All target protein signals were normalized to beta-actin signal. Then, data for the phosphorylated proteins (pSTAT3 or pERK1/2) was divided by the total protein (tSTAT3 or tERK1/2) signal data within each lane, to give a ratio for each sample. Full length, uncropped blots are presented in Supplementary Figures 6 and 7.

### sIL6Ra ELISA

The IL-6 Receptor (Soluble) Human ELISA Kit (Invitrogen; BMS214) was used to quantify soluble IL-6Ra expression in cell culture media, following the manufacturer’s instructions. For the co-culture assay (Figure 2A), media from N=4 donor lines (014_CTM, 127_CTM, M3_CTR and 069_CTF) were used. For the MGL monoculture assay to measure sIL-6Ra at timepoints through to 48h (Figure 2B), cells from the KOLF2.1J reference line (N=3 biological replicates from three distinct hiPSC-derived myeloid factories) were stimulated with acetic acid vehicle or 100ng/ml IL-6. Media was then collected after 0h (immediately after stimulation), 3h, 6h, 12h, 24h and 48h. Collected media was used undiluted and measured in duplicate on the ELISA plate to provide technical replicates. The optical density (OD) of the microwells was blanked and measured at 450nm, and referenced against 620nm. The sIL6Ra concentration was estimated against a seven-point reference curve as per the manufacturer’s instructions.

### Cytokine Profiler

Media was taken from cells in co-culture and immediately spun at 300 x *g* for 2 minutes to remove cells and cellular debris from the supernatant, which was transferred to a new tube, snap frozen and stored at -80°C until use. Co-culture media samples were grouped and pooled by treatment condition, leaving two membranes to be stained with either vehicle or IL-6 treated culture supernatant. Cytokine profiling was carried out following the manufactures instructions using the Proteome Profiler Human XL Cytokine Array Kit (R&D Systems; ARY022B). The kit contained membranes with dots of 105 different immobilised cytokine antibodies dotted with two technical replicates each (Supplementary Figure 4). Each membrane was imaged for 10 minutes using the Bio-Rad Molecular Imager^®^ Gel Doc™ XR System. Dot blot signals were quantified using the Protein Array Analyzer Palette plug-in for ImageJ, and technical dot replicates averaged to one value. These values were then corrected for background staining by subtracting the mean negative reference value from each of the signals. Each corrected value was then normalised by dividing by the mean positive reference value on the membrane.

### Custom Meso Scale Discovery Cytokine Array

Based on the previous cytokine profiler analysis, the concentrations of MIP-1a, TNF-α, IL-8, and VEGF-A in vehicle/IL-6 treated co-culture, plus 24h treated NPC and MGL monoculture media (N = 3 healthy male donors M3_CTR, 014_CTM and 127_CTM) was measured using a custom U-Plex Biomarker Group 1 (Human) Multiplex Assay (K15067M-1, Meso Scale Discovery), following the manufacturer’s instructions. Antibodies for each of the four analytes were cross-linked to a unique spot in each well of a 96-well plate. Each sample was incubated for 2 hours on the plate in technical triplicate, in addition to a seven-point standard curve. A secondary detection antibody was then added and incubated for 2 hours. The plate was read on an MESO QuickPlex SQ 120MM Imager, and data was obtained using Discovery Workbench 4.0 (Meso Scale Discovery).

### RNA extraction, RNA Library Preparation and NovaSeq Sequencing

#### RNA extraction

In total, N=4 donors were differentiated for co-culture IL-6 treatment, providing a total of 8 samples for downstream bulk RNA sequencing (RNAseq) analysis. Cells cultured for RNA extraction were collected at RT in TRI Reagent™ Solution (Invitrogen; AM9738) by splashing the cells with TRI Reagent™ Solution using a p1000, after removing media and stored at -80°C. 1ml of TRI Reagent™ Solution per 6-well was used for NPCs, and 0.5ml of TRI Reagent™ Solution per 6-well was used for MGLs. RNA was extracted from each sample by centrifugation with 200μl of 100% Chloroform (10,000 x *g* for 5 minutes at 4°C). The top aqueous layer was moved to a new 1.5ml tube with 500μl of 100% isopropanol and mixed 10 times by inversion and incubated 15 minutes at RT. Then, RNA was precipitated by centrifugation (17,000 x *g*, 15 minutes at 4°C). The supernatant was removed, and the pellet was washed in 1ml of 80% ethanol followed by centrifugation (20,000 x *g*, 5 minutes at 4°C). The ethanol was then removed, the pellet air dried for 15 minutes at RT and then dissolved in 30μl of nuclease free water. Precipitation of RNA by 0.3M Sodium-acetate and 100% ethanol at -80°C overnight was done to clean samples further, before an additional 80% ethanol wash and subsequent resuspension in 30µl RNAse-free water. Nucleic acid content was measured using NanoDrop™ One.

#### Library Preparation and NovaSeq Sequencing

Total RNA was submitted for sequencing at Genewiz Inc (South Plainfield, NJ). The following library reparations and RNA sequencing was carried out by Genewiz. Libraries were prepared using a polyA selection method using the NEBNext Ultra II RNA Library Prep Kit for Illumina following manufacturer’s instructions (NEB, Ipswich, MA, USA) and quantified using Qubit 4.0 Fluorometer (Life Technologies, Carlsbad, CA, USA). RNA integrity was checked with RNA Kit on Agilent 5300 Fragment Analyzer (Agilent Technologies, Palo Alto, CA, USA). The sequencing libraries were multiplexed and loaded on the flowcell on the Illumina NovaSeq 6000 instrument according to manufacturer’s instructions. The samples were sequenced using a 2×150 Pair-End configuration v1.5. Image analysis and base calling were conducted by the NovaSeq Control Software v1.7 on the NovaSeq instrument.

Upon receipt of FASTQ files from Genewiz, the subsequent bioinformatic analysis were performed. Files were quality controlled using Fastqc (Wingett and Andrews, 2018) and aligned to the human reference genome (GRCh38) with STAR (Dobin *et al*., 2013), then sorted and duplicates removed with samtools version 1.13 (Li *et al*., 2009) and Picard version 2.26.2 respectively, all within the King’s CREATE high performance computing systems (King’s College London, 2022). Downstream gene expression analyses were carried out in R version 4.0.2 (R Core Team, 2020). A count table was prepared and filtered for counts ≥ 1 using featureCounts (Liao *et al*., 2014) from the Rsubread (Liao, Smyth and Shi, 2019) package, version 2.4.3. Differential gene expression analysis was carried out using DESeq2 (Love, Huber and Anders, 2014) version 1.30.1 and the default Wald test. Subsequently, using the Benjamini-Hochberg (BH) method for p-value correction, only genes that passed 5% FDR were considered differentially expressed and submitted for downstream analyses.

### Over-Representation and MGEnrichment Analyses

Over representation analysis (ORA) was carried out using WebGestalt (Liao *et al*., 2019), where differentially expressed genes were tested for over representation of non-redundant cellular component, biological process and molecular function gene ontology terms. This analysis used as a background list all genes considered expressed in our model, according to DESeq2s’s internal filtering criteria (i.e., adjusted p ≠ NA). Enrichment p-values were corrected for multiple testing using the BH method, and only terms that passed 5% FDR were considered significant. To understand the similarity of the microglial gene sets generated to those from other publications, the MGEnrichment tool was used (Jao and Ciernia, 2021). Genes that were either up- or down-regulated (passing 5% FDR) were imputed into the tool, backgrounded as above. Only human pathways that were enriched were selected, and modules were considered significant if they passed 5% FDR after BH multiple testing correction.

### Immunocytochemistry

Cells were fixed with 4% paraformaldehyde (w/v; made in 4% sucrose PBS) for 20 min at room temperature. Samples were then washed twice in PBS, then simultaneously permeabilized and blocked with 2% normalised goat serum (NGS) PBS + 0.1 % Triton X-100 for 2 hours at RT. All antibodies (Supplementary Table 2) were diluted in 2% NGS in PBS. Primary antibodies were incubated with cells overnight at 4°C, and secondary antibodies were incubated with cells for 1 hour at RT, both in humidified chambers. Between incubations, cells were washed three times in PBS at 15 min intervals. Finally, cells were incubated for 10 min in PBS + DAPI (1:50,000). After the final DAPI stain, cells were left in 150µl/well PBS ready for imaging on an OperaPhenix high throughput imaging system (Perkin Elmer) using a 20x water objective over 6 to 10 consistent fields of view per well. Overall, there were 3 to 4 technical repeat wells per staining condition on each plate, coupled with 2 primary-negative wells and 2 secondary-negative wells.

#### pSTAT3/tSTAT3 Assay

Dual SMADi hiPSC-derived NPCs at D21 from three donor lines (127_CTM, 014_CTM and M3_CTR) were thawed and seeded at 25k cells per well directly into a 96-well plate for imaging. After 48h, they were treated for 24h, 3h, 30mins and 15mins with Vehicle (Acetic acid), IL-6 100ng/ml, Vehicle + rIL6Ra 100ng/ml or IL-6 + rIL6Ra. Cells were stained for TUBB3, STAT3 and phospho-(p)STAT3 and imaged as previously mentioned (Supplementary Table 2). Harmony software (Perkin Elemer) was used to identify nuclei, and the cytoplasm of each cell based on the TUBB3 stain. The raw intensity for pSTAT3 and total STAT3 was measured in both the nuclear and cytoplasmic compartments for each cell.

#### Cell-Marker Expression Assay

On day 50 of neural differentiation, 29 days after seeding at a density of 30,000 cells/well into 96-well Perkin Elmer CellCarrier Ultra plates, cells were fixed and stained. Cortical neurons were ICC stained with DAPI and TUBB3 to identify morphological boundaries, alongside glial fibrillary acidic protein (GFAP) and paired box protein (Pax6) to identify astrocytes and undifferentiated cortical NPCs respectively (Supplementary Table 2). Using the DAPI stain for PAX6 given it’s expected expression in the nucleus, and TUBB3 stain for GFAP given it’s expected expression in the cytoplasm, the number of cells positive for each marker per well was defined by the intensity of marker being above the background, which was calculated from the lowest intensity limit of each channel’s background florescence (mean 5 random locations from the primary negative wells). The number of PAX6+, MAP2+ and TUBB3+ cells per well were collated, and relative populations within each donor were calculated as a percentage of total cells counted.

#### Synapse Counting Assay

Day 50 post-mitotic cortical neurons were fixed and stained in 96-well Perkin Elmer CellCarrier Ultra plates. To identify nuclei and neurite sections on which synapses lie in cortical neurons, DAPI and MAP2 staining was performed respectively (Nieland *et al*., 2014). In addition, a combination of specific pre- and post-synaptic proteins was utilized for staining, including the pre-synaptic protein, vesicular glutamate transporter 1 (vGlut1) and post-synaptic protein, postsynaptic density protein 95 (PSD95), or Synaptic vesicle glycoprotein 2A (SV2A, pre-synaptic) and the N-methyl-D-aspartate receptor subunit GluN1(post-synaptic) (Supplementary Table 2). Furthermore, the staining also involved a combination of inhibitory synaptic proteins, Gephyrin (post-synaptic) and glutamate decarboxylase (GAD67; pre-synaptic) (Supplementary Table 2) (Shum *et al*., 2015, 2020; Deans *et al*., 2017). The following assay analysis was carried out using an automatic script, applied to all replicate wells across donor plates using the Harmony software (Perkin Elemer), with an average of 1400 cells/donor. Initially, MAP2+ cells were identified (Supplementary Figure 5). Then, dendrites were automatically identified using the MAP2+ cell masks, using a Harmony software-integrated algorithm (http://www.csiro.au) (Supplementary Figure 5). Along each dendrite, pre- and post-synaptic puncta were then identified separately from either the 568nm or 488nm imaging channel depending on the secondary antibodies used. Synaptic puncta were thresholded by two characteristics: size and intensity. Synaptic puncta sizes were considered to be between 27-170 pixels, which equated to between 0.8-5µm^2^ (Glynn and McAllister, 2006). To calculate the lowest intensity limit, the intensity of each channel’s background fluorescence was taken and averaged from 5 random locations from the primary negative wells. To identify maximum intensity, fluorescence intensity from 5 clear artefacts in the primary negative wells were averaged. The dendrite length of MAP2+ cells was measured and averaged per well, to normalise the number of puncta for culture and dendrite outgrowth. This gave two outputs for each puncta channel: intensity and puncta number/dendrite length (density) (Supplementary Figure 5). Finally, positively identified puncta were masked on top of each other, to count those that co-colocalised (50% positively overlapped). For each donor, synaptic puncta metrics were averaged from all replicate images for each condition. To reduce the possibility of confounding results caused by staining differences across donor replicate plates, all IL-6 treated puncta metrics were normalised to the vehicle average to create a fold change value from vehicle within donor.

### Statistical Analysis

All statistical analyses were conducted using Prism 9 for macOS version 9.3.1 (GraphPad Software LLC, California, USA), except for the RNAseq analyses which were carried out using the research computing facility at King’s College London, CREATE, along with R version 4.0.2 (R Core Team, 2020). Each specific test performed is described in the corresponding figure legend and supplementary statistical data table, providing the number of replicate hiPSC lines included in each technical and biological replicate, as specified in the relevant method section above. In general, when discussing co-culture data, the N=4 donors (Supplementary Table 1) were considered biological replicates, and when discussing mono-culture data the N=3 donors (3 ♂ only, Supplementary Table 1) were considered biological replicates. To compare post-mitotic cell marker populations, two-way ANOVA was used with cell type and treatment condition as main factors. When assessing the effect of IL-6 on each synaptic metric measured in post-mitotic neurons, one-sample Welch’s t-test was used due to the unequal variances between each group in the fold-change values from the vehicle-treated condition (all vehicle donor points = 1). To compare the concentrations of cytokines and sIL-6Ra in control and IL-6 treated co-culture media, an unpaired two-tailed t-test was employed. When comparing the mean between two distinct conditions, such as the concentrations of cytokines in control and IL-6 treated monoculture media from NPCs and MGLs, a two-way ANOVA was employed. For each of the cases in which a t-test was used to compare two group means, an unpaired form of t-test was used, given each cell culture sample is unique and does not reflect a paired design. In the cases where significant differences were observed between groups in these models, post-hoc testing was conducted using the Benjamini-Hochberg (BH) method with a false discovery rate (FDR) of 5% to identify the individual groups with significant differences. Adjusted p-values (q-values) less than 0.05 were considered statistically significant, and relevant significant q-values are quoted. The BH correction method with a 5% FDR threshold was also employed when deciphering the differentially expressed genes from each transcriptome signature, and during the RNAseq downstream over representation analysis.

## Results

### Forebrain neural progenitor cells only respond to IL-6 by trans-signalling in the presence of sIL-6Ra in a dose-dependent manner

To investigate whether forebrain NPCs can respond to IL-6 via trans-signalling using sIL-6Ra, we introduced recombinant human sIL-6Ra (rIL-6Ra) to forebrain NPC monocultures derived from three male donors without a psychiatric diagnosis (one clone each) at day (D) 18 of differentiation. We measured the phosphorylation of STAT3 protein (pSTAT3) levels via immunoblotting to determine if exogenous rIL-6Ra could facilitate a response to IL-6 (Figure 1A, Supplementary Figure 6). Following the combined treatment of 100ng/ml rIL-6Ra and 100ng/ml IL-6, we observed a significant increase in STAT3 phosphorylation in D18 NPCs after 15 minutes (one-way ANOVA F(3, 8) = 1773, p < 0.0001), which, while reduced, remained detectable at 3 hours (one-way ANOVA F(3, 8) = 4.920, p = 0.0318) (Figure 1B). Although SHP2 phosphorylation by JAK, which induces the Ras-ERK pathway downstream of IL-6 signaling, has been previously described in mouse immortalized fibroblasts (Fiebelkow *et al*., 2021), phosphorylation of ERK1/2 was not detected in NPCs after 15min (one-way ANOVA: F (3, 8) = 0.8869, p = 0.4881) or 3h (one-way ANOVA: F (3, 8) = 2.458, p = 0.1375) of IL-6 exposure, with or without rIL-6Ra (Supplementary Figure 7). Regarding STAT3 phosphorylation, a dose-dependent relationship was evident when varying rIL-6Ra concentrations (Figure 1C, Supplementary Figure 6). Lower doses corresponded to reduced pSTAT3/tSTAT3 ratios after 15 minutes of IL-6 exposure (two-way ANOVA: Interaction F(4, 20) = 31.94, p < 0.0001; [rIL-6Ra] Factor F(4, 20) = 36.55, p < 0.0001; Treatment Factor F(1, 20) = 436.0, p < 0.001) (Figure 1D, statistics in Supplementary Table 3). Notably, at the lowest dose (1ng/ml rIL-6Ra), there was no statistical difference between pSTAT3/tSTAT3 levels in vehicle and IL-6 treated NPCs (5% FDR = 0.371).

**Figure 1.**
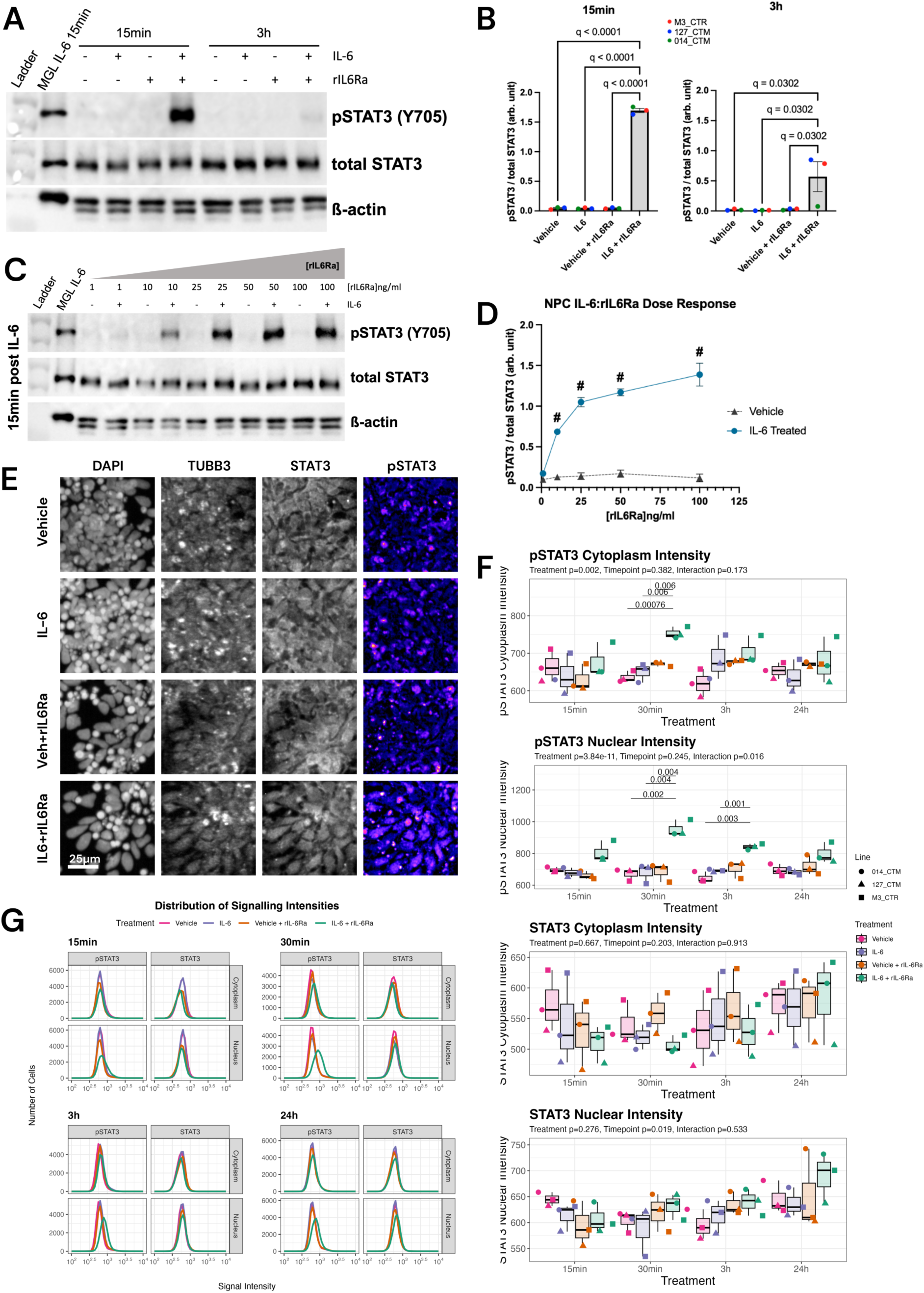
Demonstrating Trans-Signalling Capability of Cortical Neural Progenitor Cells with IL-6 in Monoculture. (A) Immunoblot depicting pSTAT3 (88kDa), tSTAT3 and β-Actin protein levels in D18 NPC monocultures. Samples, treated with vehicle or 100ng/ml IL-6 coupled with or without 100ng/ml of sIL-6Ra, were collected at 15 minutes and 3 hours for comparison. Full length, uncropped images can be found in Supplementary Figure 6. (B) Quantitative assessment of pSTAT3/tSTAT3 protein ratios from blot (A) at 15 minutes and 3 hours post-stimulation. Measurements are in arbitrary units, presented as mean ± standard deviation (SD). Statistical significance determined by one-way ANOVA with 5% FDR (Benjamini–Hochberg method) is indicated. Data derived from N=3 neurotypical male hiPSC cell lines, each with one technical replicate. Data points are colour-coded per donor line: red for M3_CTR, blue for 127_CTM, and green for 014_CTM. (C) Immunoblotting results showing 88kDa pSTAT3/tSTAT3 and β-Actin in response to a gradient of rIL-6Ra concentrations (1, 10, 25, 50, and 100 ng/ml) combined with 100ng/ml IL-6. Samples collected after 15 minutes of stimulation in D18 NPC monocultures. Full length, uncropped images can be found in Supplementary Figure 6. (D) Line graph quantification of pSTAT3/tSTAT3 protein ratios from blot (C) at 15 minutes and 3 hours, presented in arbitrary units. Data points are mean ± SD, with colour-coding indicating treatment conditions as per the key. Two-way ANOVA results, with adjustments for multiple comparisons and a q-value < 0.0001 after 5% FDR correction, are marked with ’#’ for IL-6 treatment vs. vehicle comparisons. Data based on N=3 neurotypical male hiPSC cell lines, each with one technical replicate. (E) Representative images of hiPSC-derived NPCs at D24 from donor line 127_CTM, treated for 30mins with vehicle, IL-6 100ng/ml, vehicle + rIL6Ra 100ng/ml or IL-6 + rIL6Ra. Cells were stained for TUBB3, STAT3 and phospho-STAT3. These representative images are 4x digitally zoomed from the full original 20x images available in Supplementary Figure 8. White scale bar represents 25µm. (F) Boxplots of raw staining intensities for pSTAT3 and total STAT3 in both the nuclear and cytoplasmic compartments of D24 hiPSC-derived NPCs. Each point represents the donor mean value from three technical replicate wells per condition, imaged 6 times per well. Comparison of treatment and timepoint variables were made using a two-way ANOVA, with the p-values printed above boxplots. Significant post-hoc comparisons for each sample are labelled above relevant comparison. (G) Distribution plots of signalling intensities across all cells measured across three donor lines (M3_CTR, 127_CTM and 014_CTM). Distributions are split into pSTAT3 and STAT3 stains, in both the cytoplasm and nucleus compartments of the cells. Each treatment condition is defined by a different coloured line.

**Figure 2.**
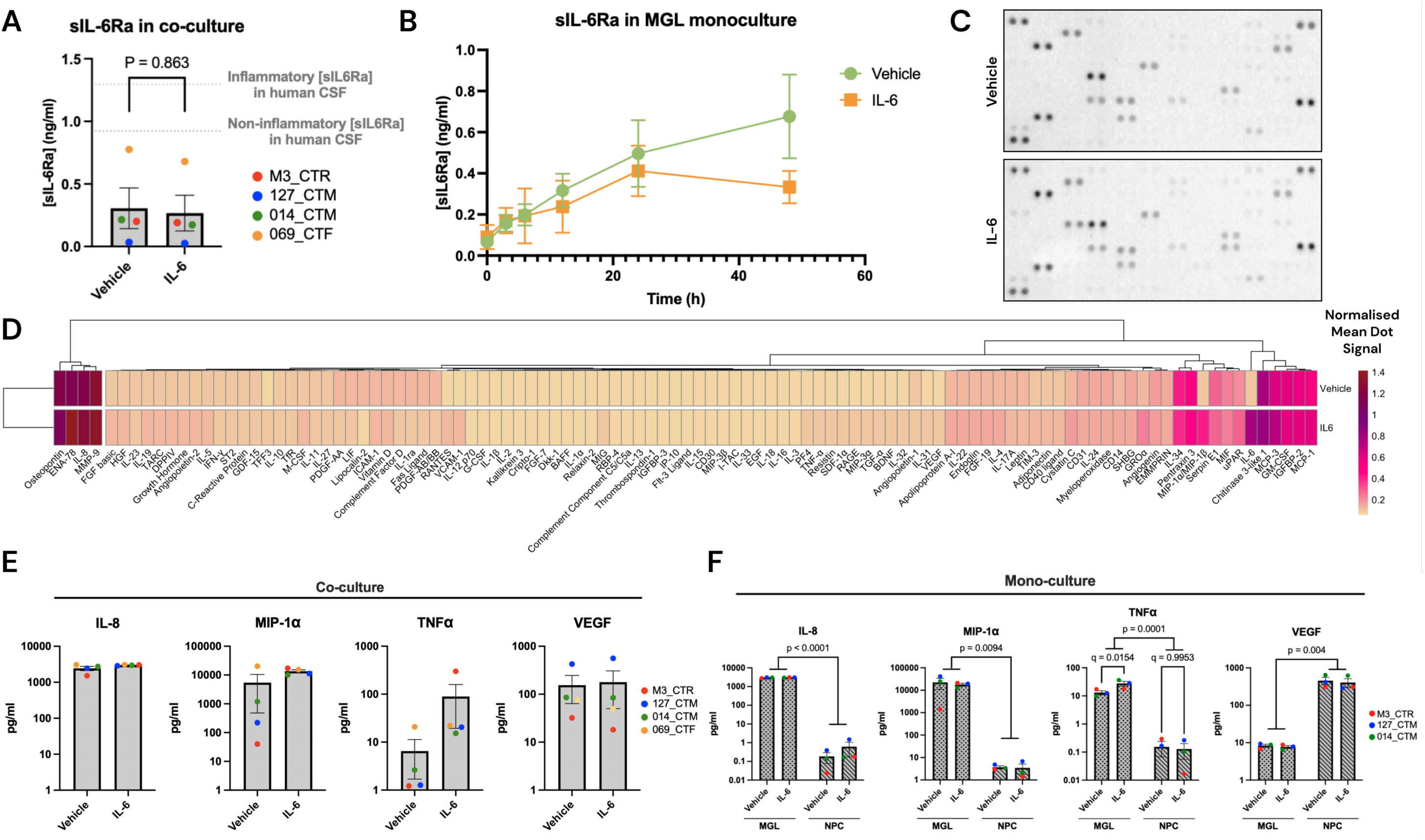
Investigating Indirect Effects of IL-6 Through the MGL Cytokine Secretion Response. (A) The sIL-6Ra concentration in co-culture media (ng/ml) varied by donor without significant IL-6 treatment effects. Data points are color-coded by donor and presented as mean ± SD. Reference lines indicate typical human CSF sIL-6Ra levels in control (non-inflammatory) and meningitis (inflammatory) cases, based on Azuma et al. (2000). (B) The sIL-6Ra concentration (ng/ml) in hiPSC-derived MGL monoculture media (N=3 myeloid factory differentiations of the KOLF2.1J line) accumulates through time to 48h but does not reach the concentration found in human CSF. (C) The cytokine profiles in vehicle and IL-6 treated NPC-MGL co-cultures as measured by dot blot images comparing cytokine secretion, using the Proteome Profiler Human XL Cytokine Array Kit (R&D Systems) measuring 104 cytokines (N = 4 donors pooled per condition). Cytokine and chemokine labels provided in supplementary material. (D) Heatmap of average raw cytokine signal, corrected for background and normalised to positive reference values per membrane, from the Proteome Profiler Array. Plotted are averaged raw signal values from 2 technical replicates per cytokine, with IL-6 added during treatment. (E) Concentrations of MIP-1a, TNF-α, IL-8, and VEGF-A measured in co-culture media after 24h with vehicle or 100ng/ml IL-6 treatment (N = 4, grey bars), as measured by the MSD multiplex cytokine assay. Data plotted as pg/ml on a log scale. Bar graphs show mean ± SD, with each point representing one donor line, and coloured as such. (F) Concentrations of MIP-1a, TNF-α, IL-8, and VEGF-A in monoculture media from control donors (N = 3 males) D14 MGLs and D18 NPCs, treated for 24h with vehicle or 100ng/ml IL-6, as measured by the MSD multiplex cytokine assay. Data plotted as pg/ml on a log scale. 5% FDR corrections post two-way ANOVA indicated. Bar graphs represent mean ± SD, color-coded by donor line.

To determine if a subpopulation of hiPSC-derived NPCs could respond to IL-6 without the presence of rIL-6Ra after 15min, 30min, 3h or 24h of stimulation, immunocytochemistry was used to stain for pSTAT3 and total STAT3 (Figure 1E, full timepoint images in Supplementary Figure 8). pSTAT3 intensity increased in both the nuclear and cytoplasmic compartments of NPCs exposed to a combined treatment of 100 ng/ml rIL-6Ra and 100 ng/ml IL-6 (Figure 1F), peaking after 30 minutes of exposure (Two-way ANOVA Cytoplasmic pSTAT3: treatment p = 0.002, timepoint = 0.382, interaction p = 0.173. Nuclear pSTAT3 intensity: treatment p = 3.84×10^-11^, timepoint p = 0.245, interaction p = 0.016). Although the intensity of total STAT3 remained unaffected by treatment in both compartments at all timepoints measured, there was an increased total STAT3 signal in all treatment groups after 24h of exposure (Two-way ANOVA Cytoplasmic STAT3: treatment p = 0.667, timepoint = 0.203, interaction p = 0.913. Nuclear STAT3 intensity: treatment p = 0.276, timepoint p = 0.019, interaction p = 0.533). When comparing the total population of cells measured in density plots, no distinct subpopulation of NPCs responded to 100 ng/ml IL-6 without rIL-6Ra, as increased pSTAT3 signal intensity shifts were only observed in cells treated with both 100 ng/ml IL-6 and 100 ng/ml rIL-6Ra (Figure 1G, cell numbers per condition in Supplementary Table 4).

### The concentration of sIL6Ra released from hiPSC-derived MGLs may not be sufficient to induce trans-signalling in NPCs

If NPCs in co-culture are to respond via trans-signalling to IL-6, then the presence of sIL-6Ra secreted by microglia is required. To confirm this, we measured sIL-6Ra concentrations in an MGL-NPC co-culture media using ELISA (Figure 2A). The results confirm the presence of the soluble receptor, but there were no significant changes in sIL-6Ra concentration after 24h of IL-6 exposure (unpaired two-tailed t-test t(6) = 0.180, R2 = 0.005, p = 0.863). The average sIL-6Ra concentration in the co-culture media was 0.29 ± 0.03 ng/ml (range: 0.27 to 0.31 ng/ml), which is lower than the average sIL-6Ra concentration typically found in human cerebrospinal fluid (CSF): approximately 1.3 ± 0.6 ng/ml in inflammatory patients (meningitis symptoms with pleocytosis) and 0.9 ± 0.5 ng/ml in non-inflammatory patients (meningitis symptoms without pleocytosis) (Azuma *et al*., 2000). Of note, the Asp358Ala A>C *IL6R* variant is known to increase sIL-6Ra secretion whilst conversely reduce a cell’s overall responsiveness to IL-6 (Ferreira et al., 2013; Khandaker et al., 2014). Therefore, we next characterized this SNP in our donor lines to mitigate potential confounding effects on our ELISA results. Genotyping revealed that all donor lines used in this study possessed the C/C allele (Supplementary Table 5). To understand the secretion dynamics of sIL-6Ra from MGLs over time, we measured the concentration of sIL-6Ra in the media after 0, 3, 6, 12, 24, and 48 hours of exposure to either vehicle or 100 ng/ml IL-6 in monoculture (N=3 independent differentiations; Figure 2B). The data confirmed an accumulation of sIL-6Ra in the media over time, with no significant effect of treatment (two-way ANOVA: treatment F(1,24) = 1.596, p = 0.219; time F(5,24) = 4.804, p = 0.0035; interaction F(5,24) = 0.797; p = 0.563). Overall, our findings indicate that although the sIL-6Ra receptor is present, the co-culture system concentration might still be insufficient for effective NPC trans-signalling during the 24-hour exposure period in the experimental set-up chosen.

Irrespective of IL-6 trans-signalling in the NPCs, we hypothesized that NPCs might additionally react to other cytokines produced by MGLs in response to IL-6 stimulation. To investigate this, we used a qualitative cytokine profiler to characterize the cytokine and chemokine milieu in the co-culture media following 24 hours of exposure to IL-6 or a vehicle control (Figure 2C). This analysis included a panel of 105 cytokines and chemokines (Supplementary Figure 4, Supplementary Table 6). We observed that osteopontin, IL-8, ENA-78, and MMP-9 were the most abundant molecules in the co-culture media across both treatment groups (Figure 5D). Relative to the vehicle control, IL-6 exposure led to increased signals in 78 molecules and a decrease in 26. The five most upregulated molecules were MIP-1α/MIP-1β (Fold Change (FC) = 4.153), RANTES (FC = 2.118), GROα (FC = 2.105), TNF-α (FC = 1.689), and PF4 (FC = 1.593). Conversely, the most significantly downregulated molecules included anti-inflammatory ligands Lipocalin-2 (FC = 0.69), PDGF-AA (FC = 0.69), LIF (FC = 0.82), and Fas Ligand (FC = 0.88).

To corroborate the findings from the proteome profiler assay, we quantitatively measured four cytokines using a multiplex cytokine assay by Meso Scale Diagnostics (Figure 2E). MIP-1α and TNF-α were selected for their upregulation in response to IL-6, while IL-8 served as a negative control due to its consistent secretion levels in both vehicle and IL-6 treated media. Additionally, VEGF was included due to its role in angiogenesis and neurodevelopment, given its observed increased expression in chronic medicated SZ patients (Chukaew *et al*., 2022) and in foetal mouse brains from an MIA rodent model (Poly I:C on gestational day 16) (Arrode-Brusés and Brusés, 2012). Consistent with our expectations, IL-8 levels remained stable with IL-6 treatment (unpaired two-tailed t-test: Fold Change (FC) = 1.22, t(6) = 1.603, R^2^ = 0.30, p = 0.160). MIP-1α and TNF-α showed an increase following IL-6 treatment, though not reaching statistical significance (MIP-1α unpaired two-tailed t-test: Fold Change (FC) = 2.48, t(6) = 1.531, R^2^ = 0.281, p = 0.177; TNF-α unpaired two-tailed t-test: FC = 13.75, t(6) = 1.180, R^2^ = 0.188, p = 0.283). In contrast, VEGF-A levels did not significantly change with IL-6 treatment (unpaired two-tailed t-test: FC = 1.155, t(6) = 0.156, R^2^ = 0.004, p = 0.884). Overall, the expression patterns of these cytokines quantitatively measured by the multiplex cytokine assay aligned with the qualitative trends observed in the proteome profiler assay, reinforcing our understanding of the cytokine response to IL-6 in our experimental setup.

Finally, to elucidate the source of cytokine secretion in our co-culture system, we examined the same cytokine candidates in 24-hour IL-6 treated D14 MGL and D18 NPC monocultures derived from N = 3 neurotypical male cultures (Figure 2F, with two-way ANOVA statistics provided in Supplementary Table 7). Our analysis indicated that MGLs were predominantly responsible for secreting IL-8, MIP-1α, and TNF-α, whereas VEGF secretion was mainly attributed to NPCs. Notably, TNF-α was the only cytokine to demonstrate a statistically significant increase in response to IL-6 treatment in MGLs (MGL Vehicle vs MGL IL-6, q = 0.015), but not in NPCs. Together, the data provide insights into the cytokine milieu downstream of IL-6 signalling in an MGL-NPC co-culture system, and the cytokines to which NPCs may respond within the co-culture environment in addition to direct IL-6 trans-signalling.

### The impact of acute IL-6 on hiPSC-derived MGL transcriptome after 24h

We next conducted bulk RNAseq analysis to characterize and confirm the overall transcriptional response of hiPSC-derived microglia-like cells (MGLs) in co-culture with neural progenitor cells (NPCs) to IL-6. Principal component analysis (PCA) of each sample highlighted the most significant variance in the MGL transcriptome: donor genetic background (PC1 = 58% variance explained) and IL-6 treatment (PC2 = 19.64% variance explained) (Figure 3A). Notably, the only female donor (069_CTF) exhibited distinct separation on the PC1 axis from male donors, but due to inclusion of a single female hiPSC line, further sex-based analysis was not pursued. Subsequent analysis revealed 72 differentially expressed genes (DEGs) post-IL-6 exposure (out of 16820 genes measured) after FDR correction at 5%. As anticipated, similarly to our prior study with IL-6 stimulated MGL mono-cultures (Couch *et al*., 2023), IL-6 receptor signal transduction genes like *JAK3* and *STAT3* were significantly upregulated (Figure 3B). The most upregulated gene was *EBI3*, known for mediating IL-6 trans-signalling, albeit less efficiently than sIL-6Ra (Chehboun *et al*., 2017). These results confirm successful IL-6 stimulation in control MGLs in co-culture.

**Figure 3.**
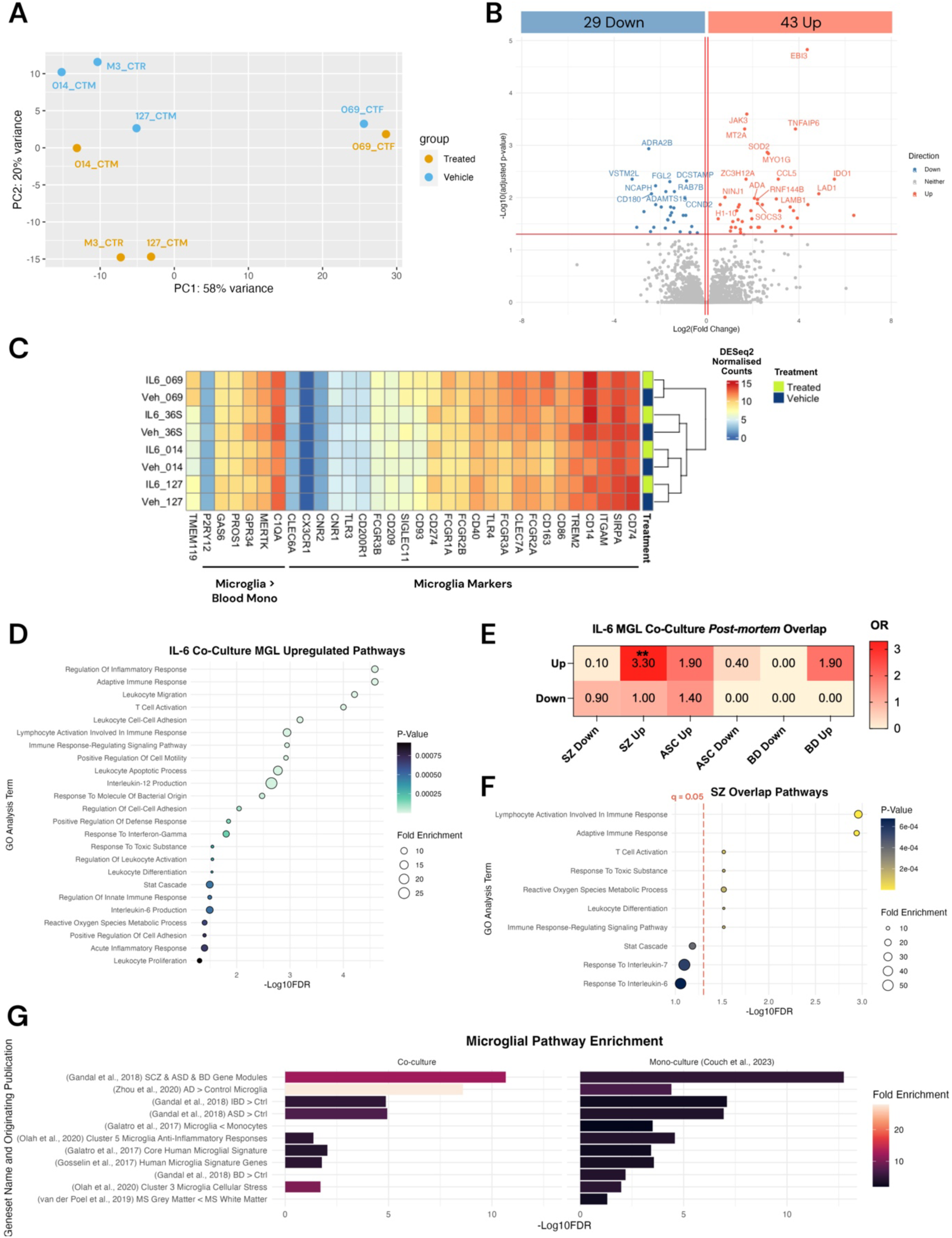
IL-6 Induces Changes in the MGL Transcriptome in Co-Culture with NPCs. (A) PCA of DESeq2 normalized counts (median of ratios), colour-coded for vehicle (blue) and IL-6 treatment (orange) and identified by donor line. Clustering reveals treatment and donor genotype influences on the MGL transcriptome. (B) Volcano plot highlighting 72 differentially expressed genes in MGLs post 24h IL-6 stimulation in co-culture with NPCs. Genes with log2FoldChange > 0.06 and adjusted p-value < 0.05 are marked in red; those with log2FoldChange < -0.06 and adjusted p-value < 0.05 in blue. Top 25 DEGs are labelled. (C) Heatmap of DESeq2 normalized counts (median of ratios) for vehicle-treated MGLs, showing consensus microglial-specific markers, including genes highly expressed in human microglia and those differentially expressed compared to blood monocytes, plus TMEM119. Right-hand side hierarchical clustering indicates no significant treatment-based changes. (D) Webgestalt gene ontology analysis of upregulated genes post-IL-6 treatment in MGLs, with 5% FDR correction. ORA terms sorted by –fadditionalFDR, colour-coded by p-value, and scaled by fold enrichment. (E) Fisher’s exact test comparing gene sets from ASC, SZ, and BD post-mortem human tissues (Gandal et al., 2018) with RNAseq-identified up- and down-regulated gene sets. Heatmap of odds ratios (OR) presented, with significance marked as: . q < 0.1, * q < 0.05, ** q < 0.01, *** q < 0.001, **** q < 0.0001; non-significant values are unlabelled. (F) Webgestalt gene ontology analysis of the 17 genes common between MGL up-regulated IL-6 and upregulated SZ post-mortem tissue gene sets (Gandal et al., 2018). Analysis is FDR-corrected at 5%, with ORA terms sorted by –log10FDR, colour-coded by p-value, and sized by fold enrichment. Red dotted line signifies q = 0.05, and values to the left of this line do not pass BH FDR correction. (G) MGEnrichment analysis of genes up-regulated in IL-6 stimulated MGLs, in both co-culture and mono-culture settings (Couch et al., 2023), adjusted with a 5% FDR. Enriched gene sets arranged by –log10FDR and colour-coded by fold enrichment.

We also validated the MGL differentiation from hiPSCs and assessed if IL-6 stimulation influenced their inherent cell type identity. The DESeq2 normalized counts of human microglia and monocyte marker genes were compared between vehicle-treated and IL-6 treated samples (Figure 3C). This comparison was based on a gene list from Haenseler et al. (2017), including genes highly expressed in human microglia (Melief *et al*., 2012) and *TMEM119*, a microglia-specific gene (Bennett *et al*., 2016). Our findings align with the expression profile of microglial marker genes as per the original MGL differentiation protocol, and no significant clustering was observed based on IL-6 treatment, suggesting that IL-6 exposure did not alter MGL cell-specificity at the gene expression level, consistent with previous data (Couch *et al*., 2023).

To explore the molecular pathways influenced by IL-6 receptor signalling in MGLs in co-culture with NPCs, we conducted a Webgestalt over-representation analysis (ORA) on the up- and down-regulated gene sets (Liao *et al*., 2019) (Figure 3D). The down-regulated gene set showed no significant associations with ORA pathways post-FDR correction (q > 0.05). In contrast, 24 pathways were significantly linked with upregulated genes, including “STAT cascade”, “Interleukin-6 production”, and “acute inflammatory response” (Figure 3D). The pathway “positive regulation of cell motility” was also notably associated, supporting previous findings that IL-6 stimulation enhances microglial motility (Ozaki *et al*., 2020; Couch *et al*., 2023). These data suggest that the observed terms related to the IL-6 pathway, along with others pertaining to cellular recruitment and cell-to-cell adhesion (such as “leukocyte migration”, “T-cell activation”, and “regulation of cell-cell adhesion”), point towards an acute, pro-inflammatory functional shift by the microglia to an acute 24h stimulation with IL-6 in co-culture with NPCs.

To understand if the IL-6-induced transcriptional changes in MGLs after 24 hours mirror those observed in neurodevelopmental disorders, we performed gene set enrichment analysis (GSEA) (Figure 3E). The analysis overlapped each signature with gene sets from post-mortem brain tissue of individuals with schizophrenia (SZ), autism spectrum conditions (ASC), and bipolar disorder (BD). Notably, the MGL transcriptome post-IL-6 exposure showed significant enrichment for the up-regulated gene set from SZ patients (list of overlap genes in Supplementary Table 8). Webgestalt ORA identified several enriched pathways, including “lymphocyte activation”, “adaptive immune response”, and “T cell activation” (Figure 3F). Collectively, these findings imply that the IL-6 response of MGLs in co-culture mirrors that observed in monocultures (Couch *et al*., 2023), and that the molecular processes identified in SZ *post-mortem* brain tissue are detectable in our developmental *in vitro* model.

Finally, we aimed to delineate the differences between the MGL transcriptomic response to IL-6 in co-culture versus mono-culture conditions (Couch et al., 2023). To achieve this, we utilised the MGEnrichment tool output to qualitatively compare specifically human modules that were enriched by up-regulated genes in both the mono-culture and co-culture MGLs following IL-6 stimulation (Figure 3G). The enrichment scores derived from our MGEnrichment analysis demonstrated a robust activation of the ’SCZ & ASD & BD Gene Modules’ in both the co-culture and mono-culture environments, indicating a highly potentiated upregulation of genes associated with NDDs. Of note, the transcriptomic profiles between mono-culture and co-culture microglia were contrasted, with a more modest NDD-related enrichment score obtained from the mono-culture microglia in comparison to the co-culture (co-culture FE = 12.36, q = 2.14×10^-11^; mono-culture FE = 5.07, q = 1.8×10^-13^). These data suggest that the presence of NPCs in a co-culture can modulate microglial gene regulation across various pathways, particularly those related to NDDs, and may suppress the MGL response to IL-6, consistent with previous findings (Haenseler *et al*., 2017).

### The proximal impact of IL-6 on hiPSC-derived forebrain NPC transcriptome in co-culture with MGLs

With the knowledge that hiPSC-derived MGLs responded to IL-6 in co-culture with NPCs, we aimed to investigate if NPCs respond to IL-6 via trans-signalling either directly from sIL6Ra secreted by MGLs, or indirectly via other cytokines and chemokines such as TNF-α secreted by MGLs in response to IL-6. Bulk RNAseq was performed on hiPSC-derived NPCs after 24 hours in co-culture with MGLs treated with either IL-6 or vehicle. We first assessed the impact of co-culturing NPCs with MGLs and IL-6 treatment on cell type heterogeneity using the NPC RNAseq dataset (Figure 4A). DESeq2 normalized counts were analysed for markers indicative of neural progenitor, pan-neuronal, dorsal forebrain, ventral forebrain, midbrain, hindbrain, upper layer and deep layer cells (Nehme *et al*., 2018) (Figure 4A). The analysis revealed predominant expression of neural progenitor cell-related genes, consistent with the expected outcome at this differentiation stage. This finding suggests that co-culturing with hiPSC-derived MGLs using trans-well inserts did not alter the anticipated NPC cell type profile, given there was no hierarchical clustering by cell-markers. Clustering was in-fact influenced by donor genotype rather than IL-6 treatment, indicating that the IL-6 exposure did not significantly affect the cell type composition of NPCs at this culture stage.

**Figure 4.**
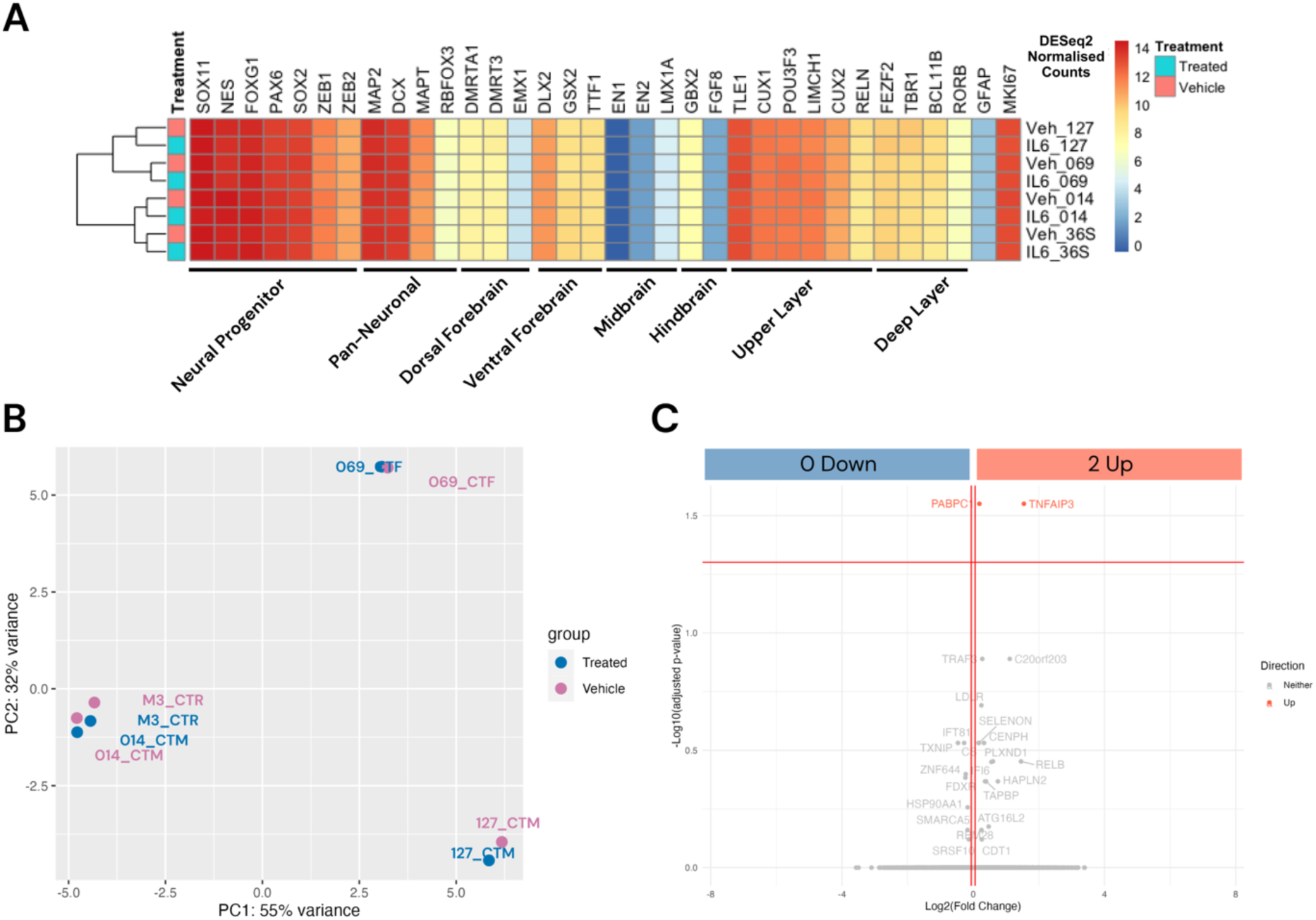
Assessing the Proximal NPC Transcriptome Response to IL-6 in Co-Culture with MGLs. (A) Heatmap of DESeq2 normalized counts (median of ratios) from vehicle-treated NPCs, displaying cell-specific marker gene expression across both genotypes. (B) PCA plot of DESeq2 normalized data (median of ratios), distinguishing vehicle (pink) and IL-6 treated (blue) samples, with donor line annotations. (C) Volcano plot showcasing NPCs’ differentially expressed genes following IL-6 exposure in co-culture with MGLs. Genes with log2FoldChange > 0.06 and adjusted p-value < 0.05 are marked in red; those with log2FoldChange < -0.06 and adjusted p-value < 0.05 in blue. Genes without a significant FDR corrected p-value are coloured grey. The top 25 expressed genes are labelled.

**Figure 5.**
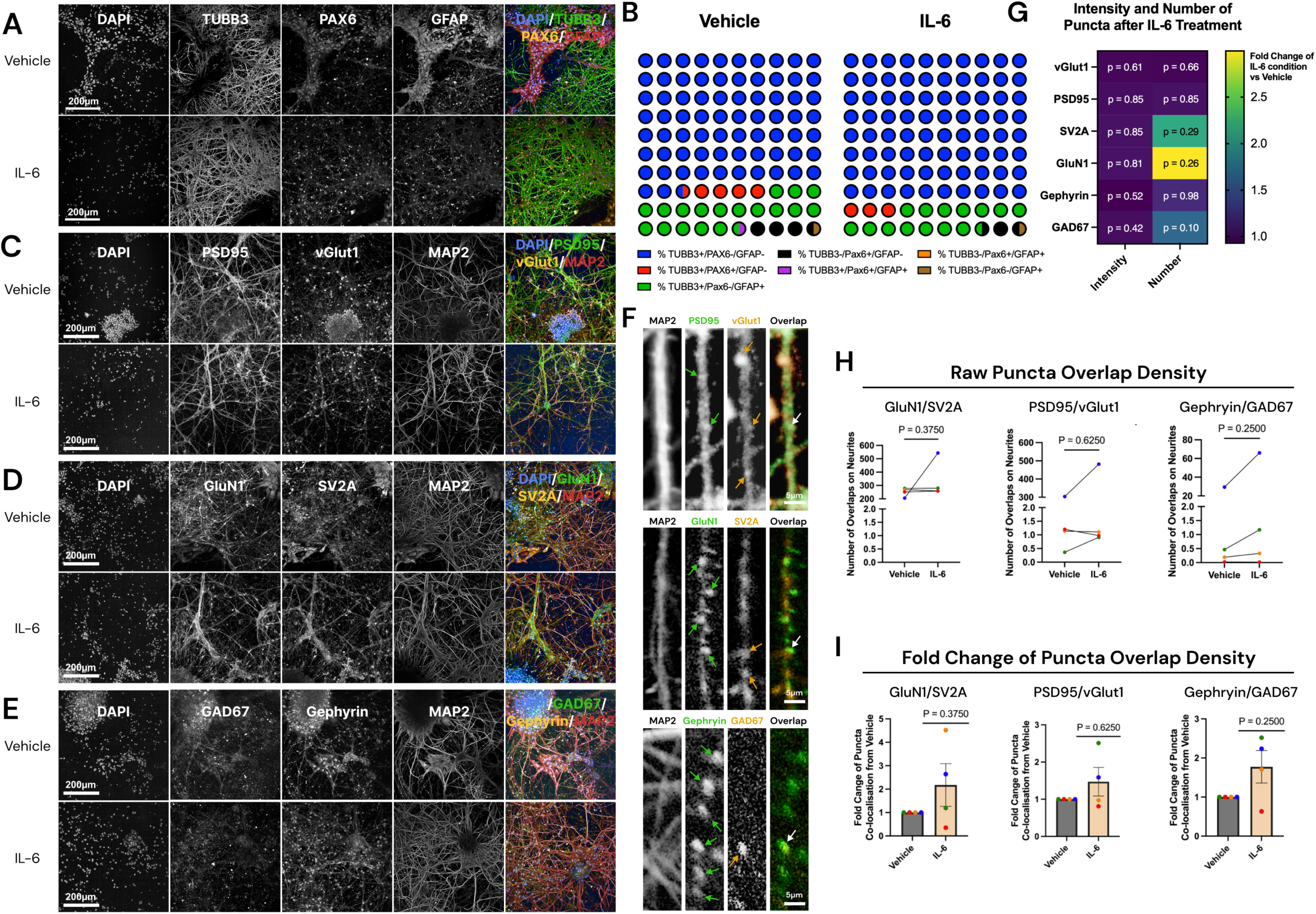
The Long-Term Effect on Terminal Neuron Differentiation After IL-6 Stimulation of NPC in Co-Culture with MGLs. (A) Representative images of cortical neurons from donor 127_CTM_01, exposed to vehicle or IL-6 in co-culture with MGLs at their NPC stage. Cells were stained for DAPI (column 1), TUBB3 (column 2), PAX6 (column 3) and GFAP (column 4). Channels were overlayed in column 5. Scale bar represents 200µm. Images from all four donors are available in Supplementary Figure 10. (B) Cell populations of the post-mitotic differentiated cultures for each treatment condition as a percentage of total number of cells counted averaged across donors (N = 4 donors averaged from N= 4 technical well culture repeats each). For clarity, percentage values were rounded to the nearest 0.5%. One circle = 1%, with corresponding percentage values found in supplementary table 9. (C-E) Representative images of cortical neurons from healthy control donor 127_CTM_01, exposed to vehicle or IL-6 in co-culture with MGLs at their NPC stage. Scale bar represents 200µm. (C) Cells were stained for DAPI (column 1), PSD95 (column 2), vGlut1 (column 3) and MAP2 (column 4). (D) Cells were stained for DAPI (column 1), GluN1 (column 2), SV2A (column 3) and MAP2 (column 4). (E) Cells were stained for DAPI (column 1), Gephyrin (column 2), GAD67 (column 3) and MAP2 (column 4). (F) Representative images of pre- and post-synaptic puncta along MAP2+ neurites from 127_CTM. Green and orange arrows indicate locations where one puncta was counted, an white arrows identify a co-localised puncta count. Scale bar represents 5µm. (G) Heatmap of fold change and p-values from one-sample Wilcoxon tests comparing vehicle (value = 1) and IL-6 treated cultures for signal intensity and number of puncta in MAP2+ neurites imaged in figures C-E. Raw metrics were averaged to give one data point per donor, and then IL-6 treated culture values were calculated as a fold change from each donor’s vehicle prior to statistical analysis. (H) Raw density of co-localisation events of puncta pairs (GAD67 with Gephyrin, GluN1 with SV2A and vGlut1 with PSD95). Wilcoxon test p-values labelled on graph compare vehicle and IL-6 treated cultures. Technical replicates were averaged to give one data point per donor. Bar graph plotted as mean with standard deviation (SD) error bars, and points coloured by donor line: red (M3_CTR), blue (127_CTM), green (014_CTM) and orange (069_CTF). (I) Fold-change from vehicle of co-localisation events of puncta pairs (GAD67 with Gephyrin, GluN1 with SV2A and vGlut1 with PSD95). One-sample Wilcoxon test p-values labelled on graph compare vehicle and IL-6 treated cultures. Similarly, metrics were averaged to give one data point per donor and then IL-6 treated culture values were calculated as a fold change from each donor’s vehicle. Bar graph plotted as mean with standard deviation (SD) error bars, and points coloured by donor line: red (M3_CTR), blue (127_CTM), green (014_CTM) and orange (069_CTF).

Contrary to our hypothesis however, PCA analysis revealed that the most significant variation in the NPC transcriptome was attributable to donor genotype rather than IL-6 treatment (Figure 4B). Out of the 17225 genes measured, only 2 DEGs passed the 5% FDR threshold for increased expression in NPCs in co-cultures treated with IL-6 versus vehicle, thereby exhibiting a minimal transcriptional difference (Figure 4C): TNF-Alpha-Induced Protein 3 (*TNFAIP3*: q = 0.0282, Log2FC = 1.545) and Poly(A) Binding Protein Cytoplasmic 1 (*PABPC1*: q = 0.0282, Log2FC = 0.1874). The absence of statistically significant changes in JAK/STAT pathway genes (*STAT3* Log2FC = 0.0997, p = 0.543, q = 0.9998; *JAK3* Log2FC = 0.584, p = 0.0396, q = 0.9998, Supplementary Figure 9) suggests that the IL-6-receptor signal transduction pathway was not activated in NPCs at the timepoint of analysis, in contrast to MGLs (Supplementary Figure 9). Additionally, the expression of four key downstream genes of the Ras-ERK pathway (*PHLDA1, DUSP4, EPHA2,* and *SPRY4*)(Chesnokov, Yadav and Chefetz, 2022), were found unchanged after IL-6 exposure in both MGLs and NPCs (Supplementary Figure 9). On the other hand, the data suggests that the NPCs might have the potential to react to the TNF-α secreted by MGLs cells, as indicated by the observed upregulation of *TNFAIP3*. In summary, under the conditions tested, forebrain cortical NPCs derived from neurotypical control donors exhibit a negligible transcriptional response to 24-hour IL-6 exposure in co-culture with MGLs.

### The long-term impact of acute NPC-stage IL-6 exposure in terminally differentiated forebrain cortical neurons

To evaluate whether IL-6 exposure to MGL-NPC co-cultures, either directly via trans-signalling or indirectly via the response secretome by MGLs, influenced the differentiation trajectories of forebrain NPCs at D18 in co-culture with MGLs, we analysed the relative proportions of cell types using cell-specific markers (Figure 5A, Supplementary Figure 10). Cell populations expressing TUBB3, GFAP and Pax6 were quantified as a percentage of total cells in terminally differentiated cortical neuron cultures at day 50 of differentiation (Figure 5B, Supplementary Table 9). The results showed that across both treatment groups, 96.66% of all cells were TUBB3+, 17.27% were GFAP+, and 6.38% were Pax6+, consistent with a predominance of neurons in the differentiated cultures. The majority of cells in both treatment groups were TUBB3+/GFAP-/Pax6-(73.43 ± 5.37%), indicating a significant proportion of neurons, followed by cells likely representing foetal astrocytes or radial glia (TUBB3+/GFAP+/Pax6-; 16.72 ± 3.04%) (Dráberová *et al*., 2008). A limited population only expressed Pax6 (TUBB3-/GFAP-/Pax6+ = 2.82 ± 1.00 %), and with minimal co-expression with TUBB3 (TUBB3+/GFAP-/Pax6+ = 3.49 ± 1.05 %) (Figure 5B). Notably, there were no significant differences in the proportions of cell types between NPCs exposed to IL-6 and those that were not (two-way ANOVA: treatment F(1,42) = 2.25×10^-7^, p = 0.999; cell-type F(6,42) = 88.04, p < 0.0001; interaction F(1=6,42) = 0.38; p = 0.89). In summary, these findings indicate that NPCs in co-culture with MGLs differentiated into a range of cell types, predominantly post-mitotic cortical neurons, and that IL-6 exposure did not significantly alter their differentiation towards various cell fates.

The architecture, number, and source of synaptic connections on neurons critically influence neuronal activity (Kuljis *et al*., 2019). Furthermore, disruptions in these synaptic properties could be central to the pathogenesis of NDDs (Bayés *et al*., 2011; Südhof, 2017; Mirabella *et al*., 2021). We therefore explored whether changes during the NPC stage, either directly by trans-IL-6 signalling or indirectly via cytokines released from IL-6 stimulated MGLs, could induce alterations in synapses. To investigate this, we quantified the density and abundance (by signal intensity) of both inhibitory and excitatory synaptic puncta along MAP2+ neurites, grown until D50 of differentiation. The choice of synaptic markers was informed by their relevance to synaptic transmission and association with neurodevelopmental disorders such as SZ and ASC. Specifically, we focused on PSD95 (Figure 5C), GluN1 (Figure 5D), and vGlut1 (Figure 5C) due to their roles in glutamatergic neurotransmission. Dysregulation in this system has been implicated in the onset of SZ and ASC (Oni-Orisan *et al*., 2008; Bitanihirwe *et al*., 2009; Moghaddam and Javitt, 2012; Holloway *et al*., 2013; De Bartolomeis *et al*., 2014; Grunwald *et al*., 2019; Mirabella *et al*., 2021). Additionally, SV2A (Figure 5D) was included due to its reported decrease in individuals with an SZ diagnosis (Onwordi *et al*., 2020, 2023). We also assessed GAD67 (Figure 5E), known for its consistently decreased expression in the brains of individuals with psychiatric disorders such as SZ, bipolar disorder (BD), and major depressive disorder (MDD) (Miyata et al., 2021; Karolewicz et al., 2010; Hashimoto et al., 2008; Guidotti et al., 2000). Lastly, Gephyrin was examined due to its association with hemizygous microdeletions in the G-domain (Figure 5E), which have been observed in unrelated individuals affected by ASC and SZ (Lionel *et al*., 2013). However, under the conditions tested, acute 24hr IL-6 exposure at the NPC stage did not significantly affect the densities, nor the abundance of either pre- or post-synaptic protein expression, as measured by the non-parametric Wilcoxon’s test (Figure 5G, Supplementary Figure 11). Furthermore, we investigated the co-localization of pre- and post-synaptic proteins: PSD95 with vGlut1, GluN1 with SV2A, and Gephyrin with GAD67 (Figure 5F). The extent of puncta co-localization, interpreted as pre- and post-synaptic connectivity, was quantified and compared between vehicle-treated neurons and those exposed to IL-6. Our results indicate that puncta co-localization was not influenced by early acute IL-6 exposure, as evidenced by the absence of significant differences between the treatment groups (Figure 5H-I). However, there was clearly extensive donor variance in these measures following exposure to IL-6, likely due individual genotype differences. Hence, we cannot conclusively dismiss the possibility of an effect of IL-6.

## Discussion

Our study aimed to determine the role of prenatal IL-6 signalling and how it might contribute to the risk of psychiatric disorders with a putative neurodevelopmental origin, by particularly focusing on the interactions between forebrain neural progenitor cells and microglia-like cells derived from hiPSCs. Our findings demonstrated that forebrain NPCs in monoculture can respond to IL-6 via trans-signalling in the presence of a recombinant soluble IL-6 receptor by activating the JAK/STAT pathway, and this response is dose-dependent on the soluble receptor concentration. This finding aligns with earlier observation that NPCs do not respond to IL-6 in monoculture in the absence of exogenous IL-6Ra or hyper-IL-6 (Couch *et al*., 2023; Sarieva, Hildebrand, *et al*., 2023). We further demonstrated that MGLs secrete soluble IL-6R and other cytokines in response to IL-6, including TNF-alpha, which has been previously identified as a key effector molecule in rodent models of maternal immune activation (Potter 2023). Contrary to our hypothesis, NPCs displayed only a marginal transcriptional response to IL-6 stimulation when in co-culture with MGLs, with only two genes showing differential expression passing the 5% FDR threshold. This finding particularly contrasts the substantial transcriptional changes observed in MGLs, indicating cell-type specific sensitivities to IL-6 signalling, as previously described in mono-culture (Couch *et al*., 2023). Moreover, concerning the enduring effects of short-term IL-6 exposure during the NPC stage, our findings revealed no marked changes in the differentiation paths of forebrain NPCs into diverse cell types. This contrasts with earlier findings in hiPSC-derived neuronal cultures that cell-fate acquisition is affected by IL-6 (Sarieva, Kagermeier, *et al*., 2023), but noting that Sarieva at colleagues (2023) stimulated their cultures with hyper-IL-6, driving trans-IL6 signalling, for five days. Furthermore, IL-6 exposure did not significantly affect synaptic densities or intensities in post-mitotic forebrain cortical neurons. Regional or species differences in the effects of IL-6 on synaptogenesis may explain why these data conflict with previous research that suggest early IL-6 exposure has the potential to alter synaptogenesis in the hippocampus of a rodent model (Samuelsson *et al*., 2006; Mirabella *et al*., 2021). It is important to emphasise that the findings of our study are of relevance primarily to dorsal and ventral forebrain NPCs. Although this study observed no specific subpopulation of NPCs to respond to IL-6 via cis-signalling, it is possible that NPCs and subsequently differentiating post-mitotic neurons from brain regions other than the developing cortex might respond differently, such as in the hippocampus or the midbrain. For example, recent evidence suggests the immortalised human dopaminergic neuronal precursor cell line LUHMES (Lotharius *et al*., 2002) expresses low levels of IL6R, albeit very low compared to microglia (Nishi *et al*., 2023). Furthermore, there are several reports in the literature on the effects of MIA in rodent models using TLR3 agonists on midbrain dopamine neuron development *in vivo*, which are associated with behavioural effects and impaired cortical function (Meyer *et al*., 2008; Vuillermot *et al*., 2010; Luchicchi *et al*., 2016; Purves-Tyson *et al*., 2021; Perez-Palomar *et al*., 2023). Therefore, we suggest our 2D hiPSC-derived forebrain co-culture model offers a simplistic, reproducible and scalable model that we can use to test hypotheses that NPCs with different regional origins, for example forebrain and midbrain, may show differential responses to IL-6 as a result.

We observe a lack of substantial transcriptional and cellular effects in NPCs co-cultured with MGLs and stimulated with IL-6, despite MGLs secreting the soluble IL-6R and other cytokines. Notably, the sIL-6Ra concentration secreted by MGLs in our study was low (0.29 ng/ml), falling below the concentration necessary to trigger STAT3 phosphorylation in NPCs in response to IL-6 and recombinant IL-6Ra (more than 1ng/ml required). Furthermore, the concentration of sIL-6Ra in our co-culture system is lower than that reported in human (adult) CSF under both basal and inflammatory disease states, acknowledging that we lack data on sIL-6Ra levels during human development in either state. Hence, there simply may not have been enough sIL-6Ra secreted by the microglia to initiate trans-signalling in the NPCs. These data highlight the importance of considering the overall human-specific cytokine milieu in our *in vitro* model, and while our iterative study initially focuses on IL-6, future research should explore species differences with other cytokines to better understand the MIA impact on foetal cortical development. Whilst mechanistic animal studies confirm that IL-6 is necessary and sufficient for many of the biological effects of maternal immune activation, the involvement of several other cytokines, including IL-1β (Allswede *et al*., 2020), IL-17A (Choi *et al*., 2016), TNF-α (Allswede *et al*., 2020; Potter *et al*., 2023), and IFN-γ (Goines *et al*., 2011; Warre-Cornish *et al*., 2020; Bhat *et al*., 2022), should not be discounted. The co-culture secretome analysis suggested that TNF-α secreted by microglia following stimulation with IL-6 could act as a potential downstream effector in NPCs. This is supported by the knowledge that NPCs possess receptors for TNF-α (*TNFRSS1A/B)* at the relevant differentiation timepoint (Couch *et al*., 2023), and can respond to this cytokine *in vivo* following maternal immune activation (Potter *et al*., 2023). Furthermore, the TNF-α family gene *TNFAIP3* was one of only two genes with differentially increased expression in the NPCs after IL-6 exposure and is known for its role in cytokine-mediated immune responses and was previously reported to be upregulated in neural stem cells infected with ZIKA virus (McGrath *et al*., 2017). The other gene with increased expression in NPCs, *PABPC1,* is involved in ribosomal recruitment and protein synthesis (Kawahara *et al*., 2008). Apart from TNF-α, the cytokines showing the greatest change in profiler signals following IL-6 stimulation in co-culture were MIP-1α/MIP-1β, RANTES, GROα, and PF4. Yet, RNAseq data from the NPCs reveal that this cell type is not likely to be the target for these cytokines, given they exhibit relatively low expression levels of specific cytokine receptors such as CCR1 and CXCR3 (normalized DESeq2 counts: 1.02 ± 0.05 and 1.85 ± 0.4, respectively) and align with those from previous forebrain transcriptomic sets (Zhang *et al*., 2016; Nowakowski *et al*., 2017; Polioudakis *et al*., 2019). Additionally, co-culturing MGLs with NPCs could induce specific cytokine responses absent in mono-cultures, as VEGF levels have been shown to be higher in co-cultures stimulated with LPS and IFN-γ (Haenseler *et al*., 2017). Future studies should investigate the impact of NPC and MGL co-culture on secreted factors from both cell types. Overall, these data highlight the significance of incorporating microglia in *in vitro* NDD models, but also cell types in addition to NPCs that can respond to the microglial secretome for a more accurate representation.

Building on this idea, only MGLs and NPCs were used in the study for simplicity, and the mechanism by which IL-6 confers increased risk for NDDs likely requires additional cell types. Consistent with this view, Campbell et al. (2014) identified cell types in mouse cerebrum and cerebellum that are responsive to IL-6 via trans-signalling, which included GFAP-positive astrocytes, Bergmann glia, lectin-bound microglia, and vascular endothelial cells, with scarce presence of IL-6 trans-signalling in neurons. Combining these rodent findings with our human cellular data suggests that IL-6 trans-signalling to neural progenitor cells and neurons may not be the direct, central pathogenic mechanism in the developing forebrain, cerebellum, and cerebrum of both species. This highlights the need for a more complex 3D model, incorporating different regional origins and additional relevant cell types, such as astrocytes, to study the effects of IL-6 on developing neurons (Haddick *et al*., 2017). Moreover, while hiPSCs provide a valuable model for studying human neurodevelopment within a human-relevant context at a molecular scale, they do not fully capture the complex interactions between additional higher-level systems seen in vivo, such as the peripheral immune system, placenta and blood-brain barrier, which would be relevant in the context of this study (Zaretsky *et al*., 2004; Campbell *et al*., 2014; Crockett *et al*., 2021; Li *et al*., 2021, 2023; Bermick *et al*., 2023). Species differences are evident in these interactions. Recent data from a mouse MIA model demonstrated that IL-6 does not pass through the placenta from the dams (Bermick *et al*., 2023), whereas IL-6 has been shown to transfer through a healthy-term human placental perfusion model (Zaretsky *et al*., 2004). It is more likely that increased maternal prenatal IL-6 triggers a larger pro-inflammatory chain reaction involving multiple cytokines, cell types, and tissues, indirectly increasing IL-6 in the foetal brain (Schaer *et al*., 2024). This limitation could be addressed by employing a human-relevant, multi-culture configuration, either in microglial-grafted organoids (Fagerlund *et al*., 2022; Pașca *et al*., 2022; Zhang *et al*., 2022; Schafer *et al*., 2023) or brain-on-a-chip fluidic systems (Liu *et al*., 2020, 2022; Vila Cuenca *et al*., 2021; Pașca *et al*., 2022). These would offer viable methods to examine whether the release of sIL-6Ra, or other cytokines and chemokines from microglia in response to IL-6 can indeed have an impact on the NPC response, via additional cell types.

Third, after only 24h of IL-6 stimulation in co-culture, cytokine secretion by MGLs in response to IL-6 response might have been either too weak or short to activate relevant pathways in NPCs. In addition, MGLs and NPCs were not in physical contact with each other, therefore precluding conclusions regarding the effects of IL-6 that may occur in a physical co-culture system due to cell-to-cell contact. Our trans-well co-culture model was designed to test hypotheses relating to secreted factors from the MGLs and avoid potential confounds related to cell sorting (Cadiz *et al*., 2022; Akhmetzyanova, Rizvanov and Mukhamedshina, 2024) in identifying transcriptional effects of IL-6 on MGLs and NPCs in this system. The complexity of a chain of signalling pathways from additional cell types or physical cell connections over a more chronic period may be required to provoke an IL-6 effect on NPC development, which our *in vitro* setting may not entirely replicate, but could be addressed with studies using 2D and 3D cultures in combination with single-cell sequencing. Furthermore, NPCs underwent passaging following IL-6 exposure, potentially biasing the sample towards healthier NPCs, leaving behind the ones most affected by IL-6. Therefore, it is important to consider that IL-6 exposure may have long-term effects on developing NPCs when in contact with hiPSC-derived MGLs, and not subjected to post-stimulation passaging. As a result, our study suggests that physical 2D culture and a more prolonged cytokine exposure remains important to test in order to induce substantial changes in NPCs, such as the 5-day exposure to brain organoids by Sarieva and colleagues (2023). Such extended exposure and longer-term culture could more closely mimic the physiological conditions of a long-term prenatal infection, potentially offering a more realistic model for studying the effects of maternal immune activation on foetal brain development (Sarieva, Kagermeier, *et al*., 2023).

Fourth, the noticeable presence of variation between different donor lines may mask detectable differences at an averaged donor level. Our sample size of N=4 donor lines, although in line with recommendations for the minimum sample size in such experiments (Dutan Polit *et al*., 2023), coupled with the variability in responses generated by each individual donor may well constrain our ability to detect group-level differences. Additionally, the diversity among our donor samples may be influenced by sex, as illustrated in Figure 3A PCA, where the sole female line (069_CTF) separates from the three male lines along PC1. The inclusion of only one female line however precludes any conclusion with regard to potential differences in the response to IL-6 stimulation as a function of chromosomal sex (XX vs. XY). Future studies using equivalent numbers of XX and XY donor lines or designs incorporating isogenic lines with different sex chromosome content will be required to address this question (Waldhorn *et al*., 2022). Taken together, these overall sources of variation are consistent with the known heterogeneity of outcomes in response to maternal immune activation in humans and in rodent models, and that in most cases there is no negative outcome in the offspring (Meyer, 2019; Mueller *et al*., 2019). Therefore, the interaction between environment and genetic background needs to be considered, as evidenced by our findings of differential transcriptional response of NPCs from individuals with schizophrenia to IFN-γ stimulation (Bhat *et al*., 2022). Given the polygenic nature of NDDs and psychiatric disorders, we suggest hiPSC models that retain the genetic background of the donor are ideally placed to test this (Jansen *et al*., 2020; Singh *et al*., 2022; Trubetskoy *et al*., 2022). When conducting studies to examine the roles of cytokines on human neurodevelopment *in vitro* using healthy donor lines however, either greater statistical power, or stratification to allow selection of specific individuals to study are critical factors to consider when designing future studies. As such our data may be thought of as part of an iterative process to refine the use of hiPSC models to study how cytokines influence neurodevelopment in health and disease states.

Despite the minimal impact of IL-6 stimulation on NPCs within our co-culture system there are clear and replicable effects of IL-6 on hiPSC-derived MGLs that are of relevance and open new avenues for research. For example, data from mice suggests that MIA accelerates the developmental programming of microglia, impairing their intended developmental functions (Matcovitch-Natan *et al*., 2016). Our previous work demonstrated that the “MIA Poly I:C GD14 P0” module from Matcovitch-Natan et al. (2016), a microglial gene set from newborn pups exposed to Poly I:C on gestational day 14, was the most relevant module overlapping with our upregulated gene set in hiPSC-derived MGL monocultures exposed to IL-6 for 3h (Couch *et al*., 2023). Additionally, a recent study that analysed transcriptomes and epigenetic profiles at various developmental stages in both foetal and postnatal human microglia has identified IL-6 as ligand of particular significance in predicting the postnatal microglial transcriptome, indicating that IL-6 may play a role in promoting the development of a mature, postnatal microglial phenotype (Han *et al*., 2023). This raises the idea that the primary mechanism by which IL-6 increases NDD risk prenatally in offspring could be via the alteration of typical developmental functions of microglia following IL-6 exposure. This guides several important questions unaddressed by the present study; are the MGLs in this study are responding to IL-6 through cis- or trans-signalling what causes the functional shift between these two signalling states, and (3) whether and how this signalling state transition operates in the CNS, all of which need to be addressed in future studies. Distinguishing these effects can be carried out in specific studies to block the IL-6 trans-signalling pathway in microglia using a soluble IL-6ST compound and compare functional outcomes. Overall, these aspects represent a critical area for further investigation to understand the complex dynamics of neurodevelopmental changes under maternal immune activation and the influence of IL-6 on NDD risk.

In conclusion, our findings show that hiPSC-derived NPCs respond by trans-signalling to IL-6 when sIL-6Ra is present, above a concentration of 1ng/ml, a level not met by hiPSC-derived MGLs from healthy donors *in vitro* and unchanged by acute IL-6 stimulation. Although these MGLs secrete a wide range of other cytokines and chemokines including TNF-α in response to IL-6, the proximal transcriptional and longer-term cellular effects on NPCs were minimal to absent. On the other hand, MGLs in co-culture showed robust transcriptional responses to IL-6 consistent with our prior observations and raising interesting new questions. Based on our observations we suggest that future studies seeking to develop *in vitro* hiPSC models to decipher the effects of IL-6 on human neurodevelopment should focus on incorporating microglia into more complex 3D models containing all relevant glial and neuronal cells that may respond to the microglia secretome. Such experiments should ideally be carried out on appropriate genetic risk backgrounds such as high polygenic risk for schizophrenia, which may reveal important disease-relevant mechanisms. Alternatively individual donors could be selected based on multivariate data from deeply-phenotype birth cohorts to identify cellular and molecular mechanisms associated with risk or resilience to IL-6 exposure during neurodevelopment.

## Acknowledgements

The authors acknowledge use of King’s Computational Research, Engineering and Technology Environment (CREATE) and are grateful to Dr George Chennell of the Wohl Cellular Imaging Centre at King’s College London for technical support during live imaging. ACMC, DPS and ACV acknowledge financial support for this study from the National Centre for the Replacement, Refinement and Reduction of Animals in Research (NC/S001506/1). RM and AB are in receipt of the MRC-Sackler Ph.D. Programme studentship as part of the MRC Centre for Neurodevelopmental Disorders (Medical Research Council MR/P502108/1). LS is supported by the UK Medical Research Council (MR/N013700/1) and King’s College London member of the MRC Doctoral Training Partnership in Biomedical Sciences. CR and ACV acknowledge support by the Neuro-Immune Interactions in Health & Disease Wellcome Trust PhD Training Programme (218452/Z/19/Z) at King’s College London.

