## Supplementary File for "Transcriptional and Cellular Response of hiPSC-derived Microglia-Neural Progenitor Co-Cultures Exposed to IL-6"

***gDNA extraction, PCR, Gel Electrophoresis and Sanger Sequencing***

**gDNA Extraction:** the Monarch® Genomic DNA Purification Kit (New England BioLabs; #T3010S/L) was used to extract genomic (g)DNA from frozen cell pellets, originating either from old myeloid factories or hiPSC cultures. Frozen pellets were thawed slowly on ice and resuspended in 100µl cold PBS. 1µl Proteinase K and 3µl RNase A was added to the resuspended pellet and mixed, before 100µl Cell Lysis Buffer was added and vortexed. The solution was then incubated for 5mins at 56ºC with agitation. After incubation, 400µl of gDNA Binding Buffer was added to the lysate and vortexed. The entire resulting lysate was added to the gDNA Purification Columns and centrifuged for 3mins at 1000 x *g* to load the column and then for 1min at 12,000 x *g* to clear the column. The column was then transferred to a new collection tube and washed twice with 500µl gDNA Washer Buffer for 1min at 12,000 x *g* and flowthrough discarded. Finally, gDNA was eluted into a fresh Eppendorf tube after adding 50µl preheated (60ºC) gDNA Elution Buffer. Nucleic acid content and quality was measured using NanoDrop™ One.

**Polymerase Chain Reaction:** Amplification of extracted gDNA was performed by PCR. Two distinct primers were designed to flank the *IL6Ra* target region of interest (Fw: AGGAGGTCCCCAAAGCCTTCCG, Rv: GGCGAGATTGCACCACAGCACT) with an optimal annealing temperature of 72ºC using the online browser Benchling (www.benchling.com). In a total volume of 50µl, 100ng of gDNA was added to 25µl Q5 Hot Start High-Fidelity 2x Master Mix (New England BioLabs; #M0494S) and 1µM of both forward and reverse primer, and cycled in conditions as follows: initial denaturation at 98°C for 30 seconds, followed by 35 cycles of denaturation at 95°C for

10 seconds, annealing at 72°C for 30 seconds, and extension at 72°C for 30 seconds, with a final extension step at 72°C for 10 minutes.

**Agarose gel electrophoresis:** to visualise PCR products, a 1.5% agarose gel was made by diluting 1.5 g of agarose powder in warm 100ml 1X TAE. Once the agarose was dissolved and the solution was clear, it was slightly cooled under a cold-water tap. Only once cool was ethidium bromide (0.002%) added to the 1% agarose and the gel was poured and left to set. Once set, it was placed in a running tank and covered with 1X TAE buffer. 5µl of PCR samples were loaded to their designated wells along with 5 µL of Quick-Load® 1 kb Plus DNA Ladder (New England BioLabs; N0469S). Samples were loaded at the negative cathode end of the electrophoresis, since when a current is applied DNA will move towards a positive anode given its inherent negative charge owing to its phosphate backbone. The samples were run at 100V for 30 minutes, and the gel visualized under a UV gel visualizer.

**Sanger Sequencing:** We verified the specific variations of the IL6Ra Asp358Ala variant in each donor line using Sanger sequencing. For this purpose, we sent 5µl of PCR products along with 5µl of forward primers (at a final concentration of 0.3µM) to Eurofins Genomics LLC and Source BioScience for sequencing. The resulting Sanger sequences were analysed using the Snap Gene application (version 6.2.0) to confirm the genotypes. Screenshots of these sequences have been included for reference in Supplementary Table 5.

**Supplementary Table 1 -** Donor lines used throughout study.

| Donor | Line Clone | Genotype | Reprogramming | Cohort | Age | Sex |
|---|---|---|---|---|---|---|
| **014_CTM** | 014_CTM_02 | Control | CytoTune™ Sendai | LEAP / StemBANCC | 18-30 | Male |
| **M3_CTM** | M3_CTM_36S | Control | CytoTune™ Sendai | EU-AIMS | 18-30 | Male |
| **127_CTM** | 127_CTM_01 | Control | CytoTune™ Sendai | EU-AIMS | 51 - 60 | Male |
| **069_CTF** | 069_ CTF_01 | Control | CytoTune™ Sendai | EUAIMS / StemBANCC | 31-50 | Female |

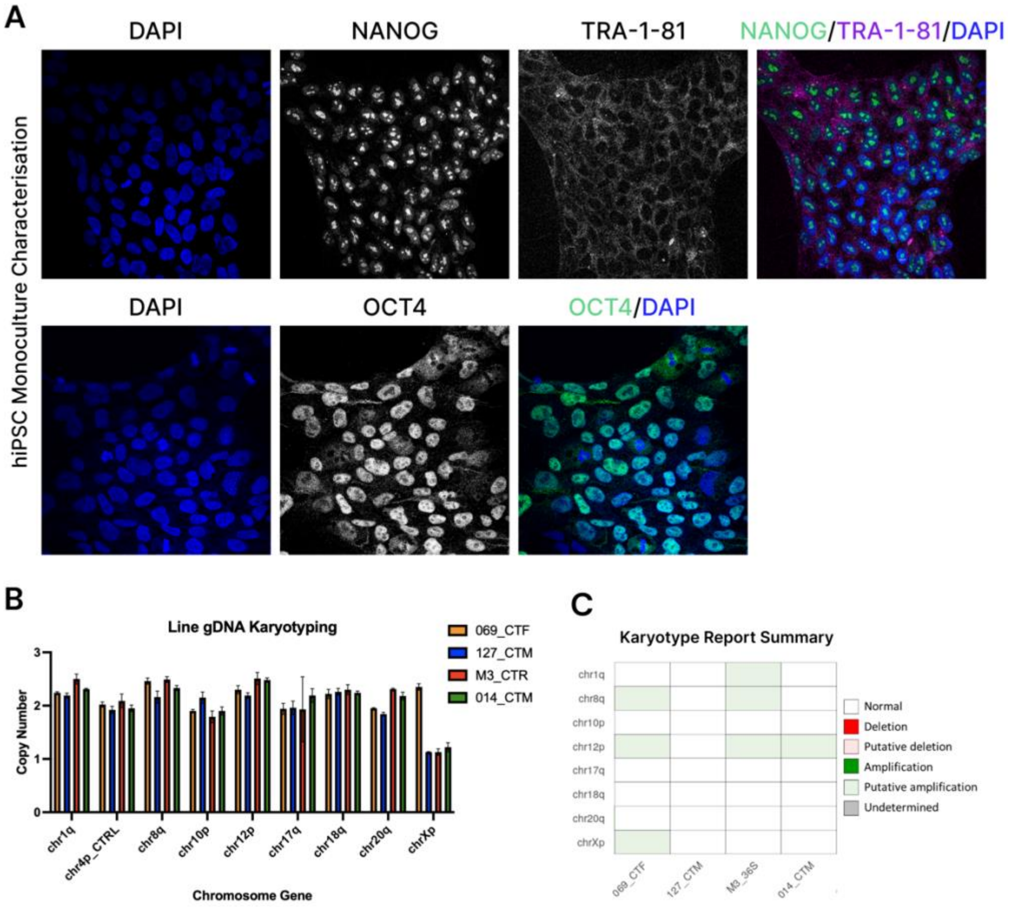


**Supplementary Figure 1 – hiPSC line characterisation.** (A) hiPSCs were stained for pluripotency markers Nanog (Abcam #80892) , OCT4 (Invitrogen #701756) and TRA-1-81 (Invitrogen #MA1-024), as per the ISSCR guidelines. Images from M3_CTR are represented in this figure. (B) gDNA from the lines used to generate the NPC and MGL cultures were also karyotyped using the hPSC Genetic Analysis Kit from StemCell Technologies (#07550), following the manufacturers protocol. Copy number variants for karyotypic abnormalities reported in human hiPSC lines are displayed in the bar chart, coloured by cell line. (C) The heatmap summary of copy number variants for each line identified no significant abnormalities in the lines used in this study, other than putative amplifications at: chrXp, chr12p and chr8q for the 069_CTF line; chr12p, chr8q and chr1q for the M3_CTR line; and chr12p for the 014_CTM line.

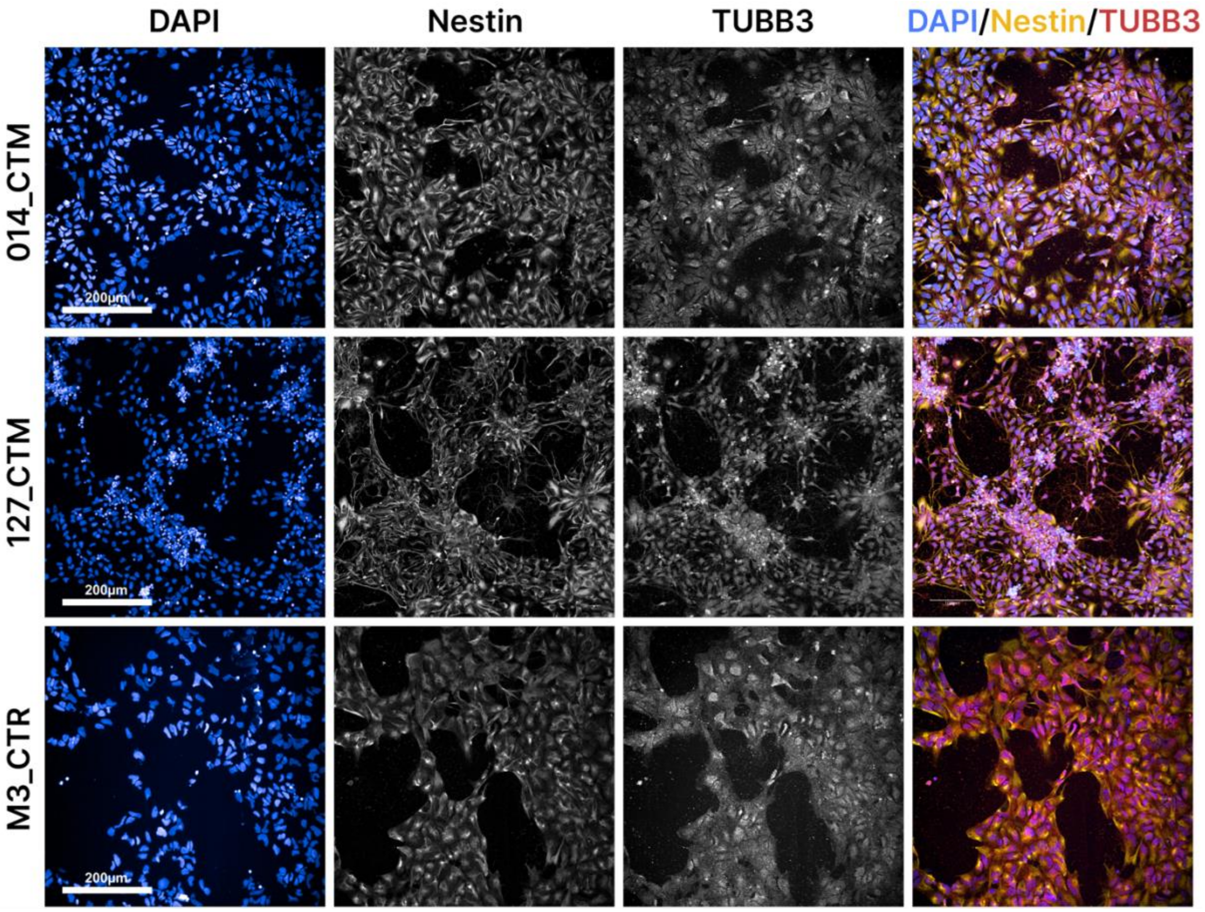


**Supplementary Figure 2 – NPC quality control immunocytochemistry**. Dual SMADi hiPSC_derived NPCs were generated from three donor lines (014_CTM, 127_CTM and M3_CTR) and stained for typical neural progenitor marker Nestin (R&D systems Md #MAB1259 1:500) and neuronal marker TUBB3 (Aves labs Rb #TUJ) to confirm cellular identity. Scale bar represents 200µm.

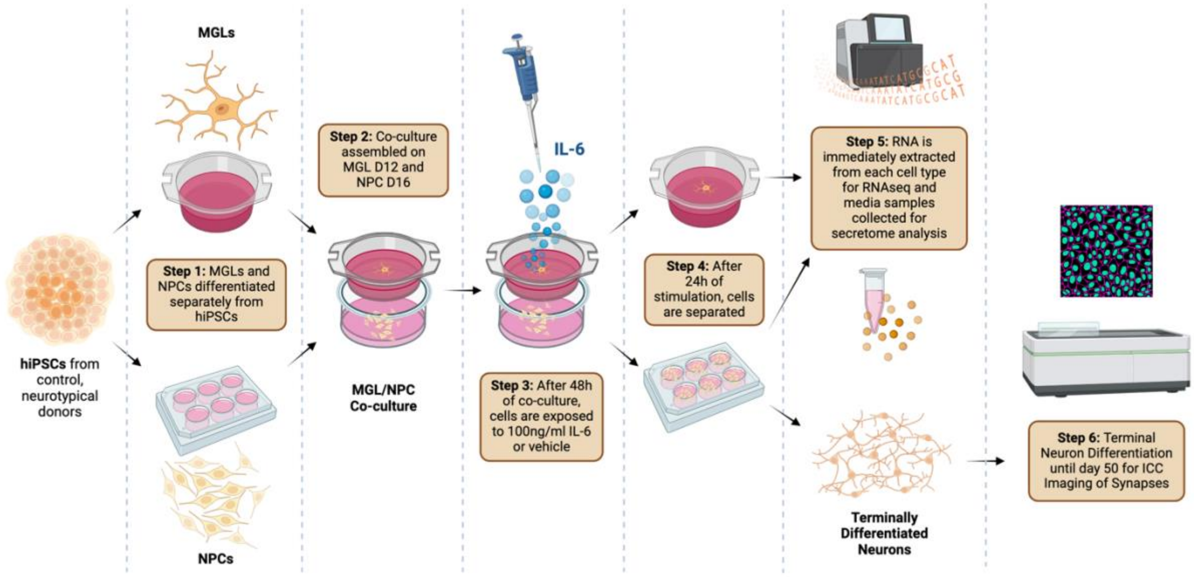


**Supplementary Figure 3 - Scheme for exposing co-cultured hiPSC-derived neural progenitor cells (NPCs) and microglial like cells (MGLs) with 100ng/ml IL-6.** Day 12 MGLs and day 16 NPCs were derived separately on cell culture inserts and 6-well plates respectively (step 1). They were then co-cultured for 48h (step 2) before stimulation with 100ng/ml IL-6 or acetic acid vehicle for 24h (step 3). MGL and NPC RNA and media samples were then collected after compartment separation (step 4) and extracted for analysis (step 5).

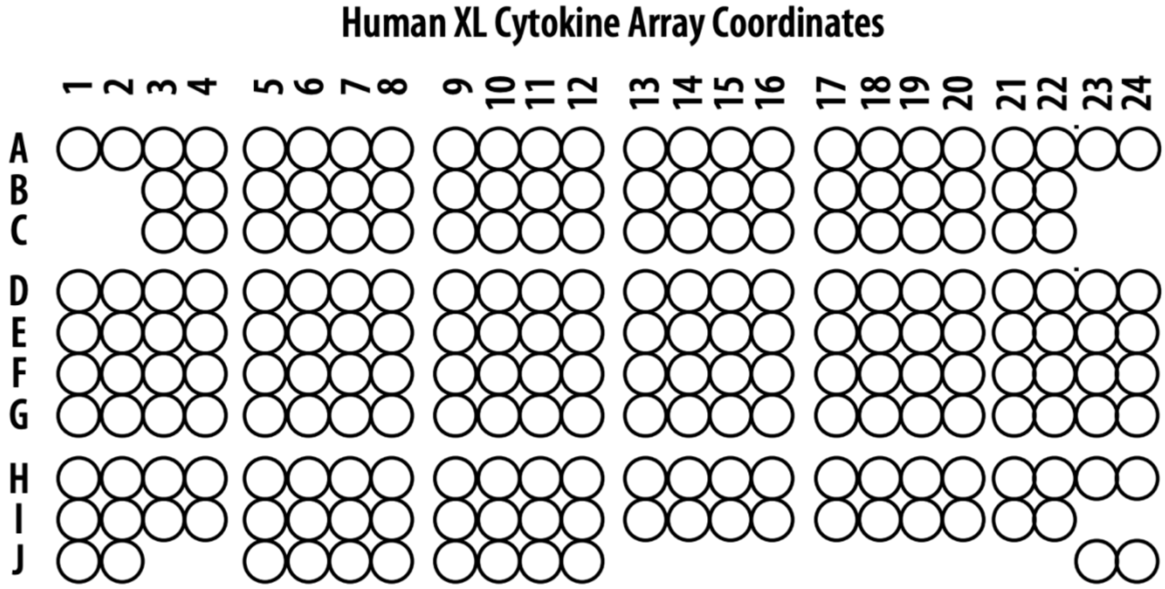


**Supplementary Figure 4 -** Human cytokine array layout on each membrane of the small, 36 cytokine Proteome Profiler Human XL Cytokine Array kit from R&D systems.

**Supplementary Table 2 - List of antibodies used for immunocytochemistry assays.**

| Assay | Epitope | Host | Dilution | Supplier |
|---|---|---|---|---|
| **pSTAT3/tSTAT3** | TUJ1 (TUBB3) | Chicken | 1:1000 | Aves labs; TUJ |
| | STAT3 | Mouse | 1:800 | Cell Signalling; 9139 |
| | Phosphorylated-STAT3 | Rabbit | 1:100 | Cell Signalling; 9145 |
| | Anti-rabbit AlexaFluor 488 | Goat | 1:750 | Thermo Fischer Scientific; A11098 |
| | Anti-mouse AlexaFluor 568 | Goat | 1:750 | Thermo Fischer Scientific; A11004 |
| **Synapse Counting Assay** | MAP2 | Chicken | 1:1000 | Abcam; ab92434 |
| | GluN1 (NMDAR1) | Mouse | 1:500 | Biolegend; 818601 |
| | vGlut1 | Rabbit | 1:500 | Synaptic Systems; 135302 |
| | PSD95 | Mouse | 1:500 | Biolegend; 810401 |
| | Gephyrin | Rabbit | 1:500 | Millipore; AB5725 |
| | GAD67 | Mouse | 1:500 | Abcam; ab26116 |
| | SV2A | Rabbit | 1:500 | Abcam; ab32942 |
| **Cell-Marker Expression Assay** | GFAP | Chicken | 1:500 | Abcam; ab4674 |
| | TUJ1 (TUBB3) | Mouse | 1:500 | Biolegend; 801201 |
| | Pax6 | Rabbit | 1:500 | Biolegend; 901301 |
| **Cell-Marker Expression and Synapse Counting Assays** | Anti-rabbit AlexaFluor 568 | Goat | 1:750 | Thermo Fischer Scientific; A11011 |
| | Anti-mouse AlexaFluor 488 | Goat | 1:750 | Thermo Fischer Scientific; A11001 |
| **pSTAT3/tSTAT3, Cell-Marker Expression and Synpase Counting Assays** | Anti-chicken AlexaFluor 633 | Goat | 1:750 | Thermo Fischer Scientific; A21103 |

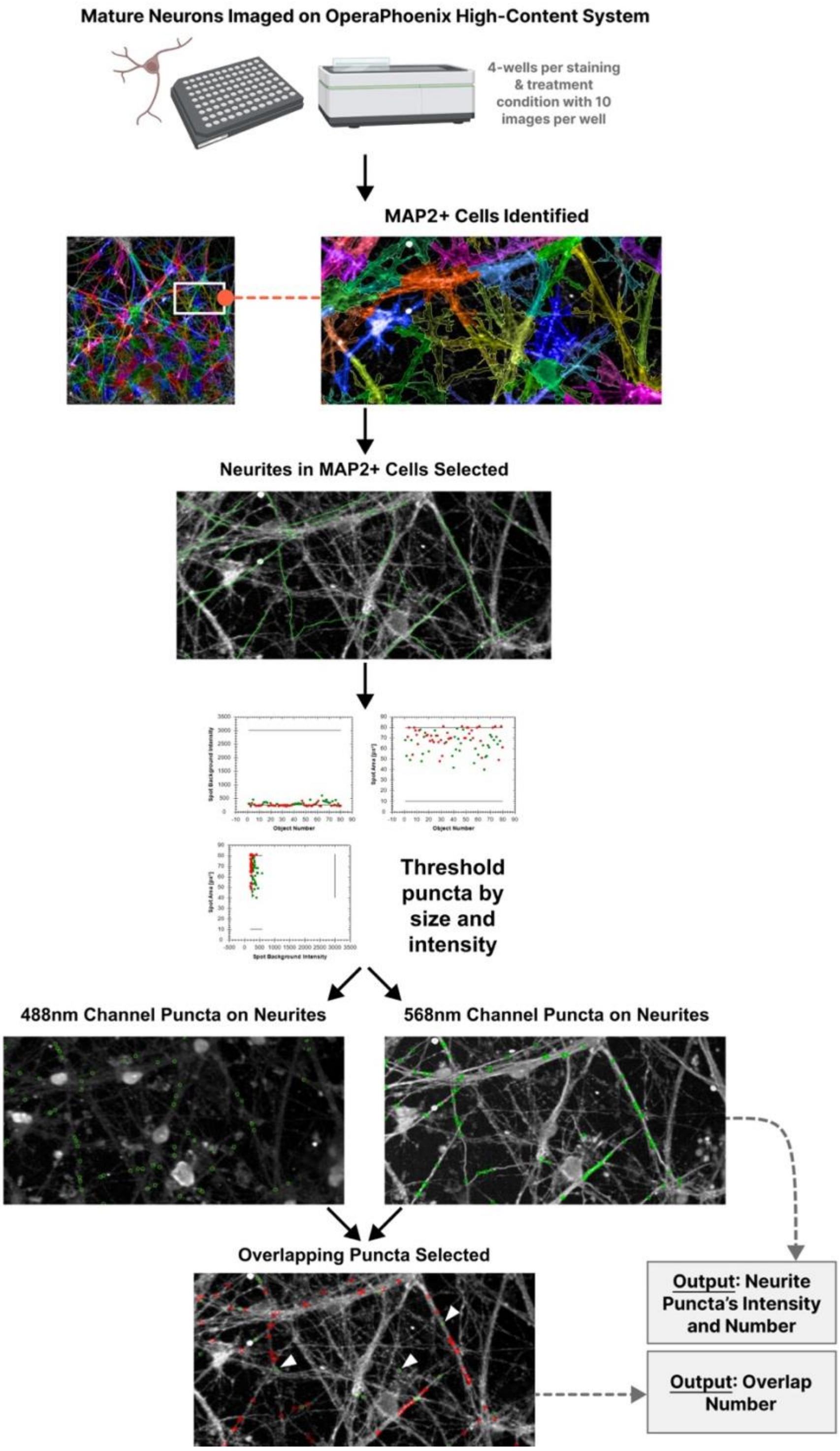


**Supplementary Figure 5 - Schematic for Synapse Counting.** Mature cortical neurons that had been exposed to acute IL-6 treatment in co-culture with MGLs at an NPC stage were seeded into a 96-well plate. MAP2+, DAPI and combinations of synaptic puncta were imaged on the OperaPhoenix high-content screening system. Following this, images were analysed on Harmony software to identify Map2+ cells, and subsequent neurites indicated by green lines. Puncta along these neurites were found by thresholding between size and intensity limits. Positively identified puncta were indicated by the software with a green circle, counted, and their fluorescence intensity measured. Finally, co-localisation of puncta in both channels were counted, with positive co-localisation targets identified with a green circle and white arrow.

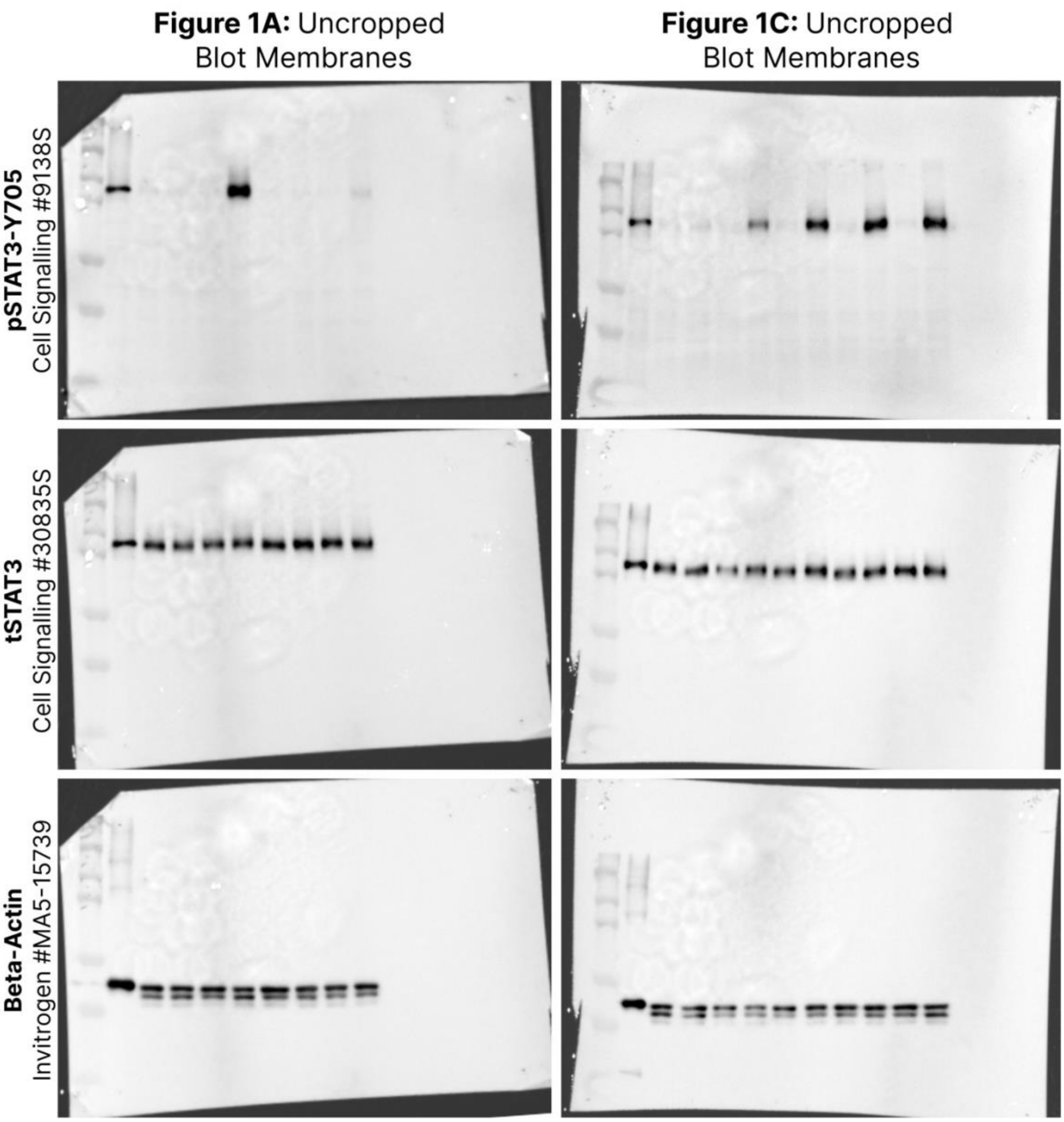


**Lane map: (1)** Dual Color standards marker (BioRad;1610374), **(2)** 15min MGL IL-6 positive control, **(3)** 15min Vehicle, **(4)** 15min 100ng/ml IL-6, **(5)** 15min Vehicle + 100ng/ml rIL6Ra, **(6)** 15min IL-6 100ng/ml + 100ng/ml rIL-6Ra, **(7)** 3h Vehicle, **(8)** 3h 100ng/ml IL-6, **(9)** 3h Vehicle + 100ng/ml rIL6Ra, **(10)** 3h IL-6 100ng/ml + 100ng/ml rIL-6Ra.

**Lane map: (1)** Dual Color standards marker (BioRad;1610374), **(2)** 15min MGL IL-6 positive control, **(3)** 15min Vehicle + 1ng/ml rIL6Ra, **(4)** 15min IL-6 100ng/ml + 1ng/ml rIL-6Ra, **(5)** 15min Vehicle + 10ng/ml rIL6Ra, **(6)** 15min IL-6 100ng/ml + 10ng/ml rIL-6Ra, **(7)** 15min Vehicle + 25ng/ml rIL6Ra, **(8)** 15min IL-6 100ng/ml + 25ng/ml rIL-6Ra, **(9)** 15min Vehicle + 50ng/ml rIL6Ra, **(10)** 15min IL-6 100ng/ml + 50ng/ml rIL-6Ra, **(11)** 15min Vehicle + 100ng/ml rIL6Ra, **(12)** 15min IL-6 100ng/ml + 100ng/ml rIL-6Ra.

**Supplementary Figure 6 –** Uncropped, full-length blots that correspond to figures 1A (column 1) and 1C (column 2), to demonstrate antibody specificity. Blots were probed in order of pSTAT3, tSTAT3 and subsequently beta-actin. Blots were stripped between each primary antibody probe; hence, identical membranes are presented in each column of this figure. A lane map identifying each lane's sample is written below each membrane.

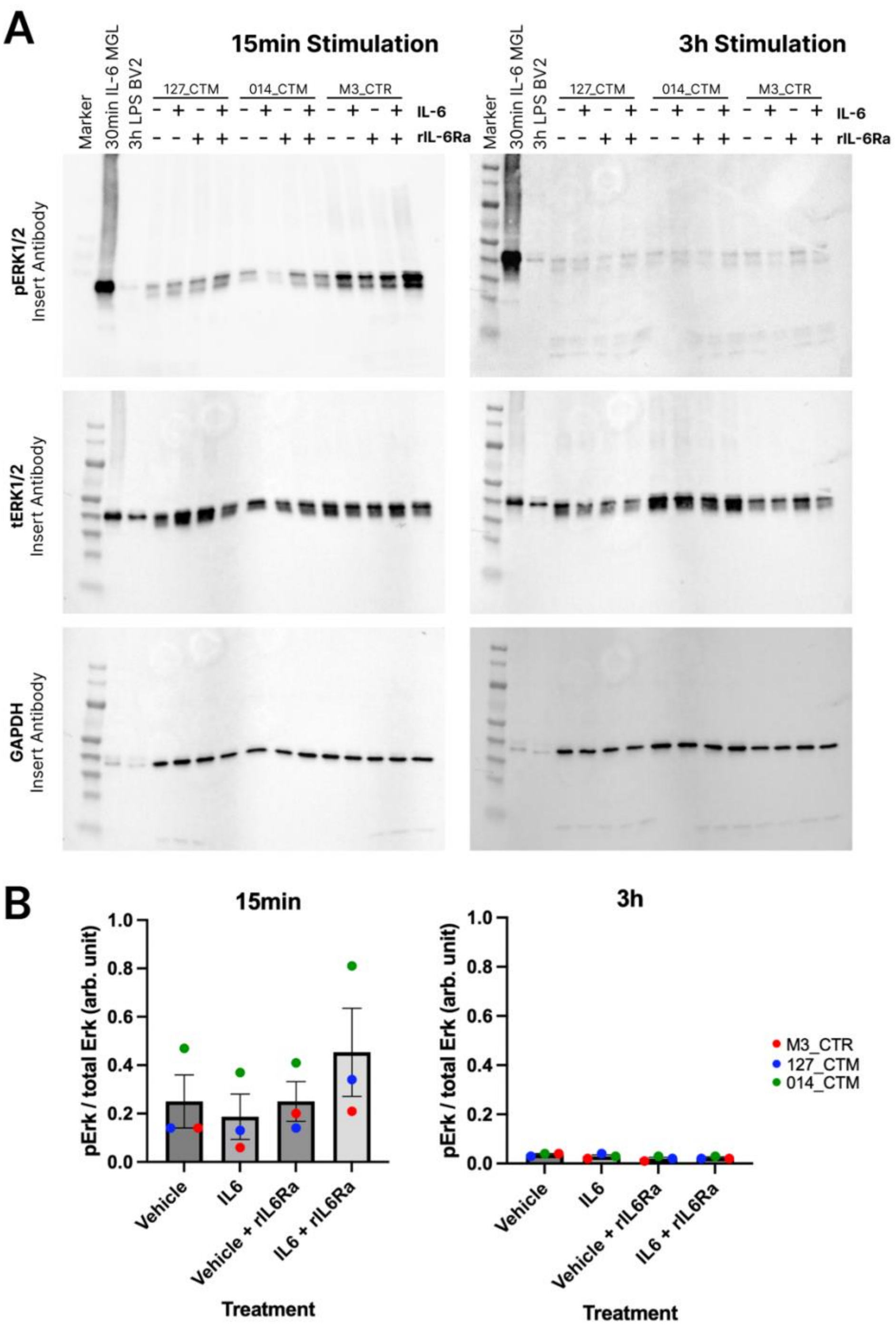


**Supplementary Figure 7 –** hiPSC-derived NPCs do not activate the Ras-ERK pathway after 15min or 3h IL-6 exposure. (A) Immunoblot depicting pERK1/2, tERK1/2 and GAPDH protein levels in N=3 (M3_CTR, 127_CTM and 014_CTM) D18 NPC monocultures. Samples, treated with vehicle or 100ng/ml IL-6 coupled with or without 100ng/ml of sIL-6Ra, were collected at 15 minutes and 3 hours for comparison. (B) Quantitative assessment of pSTAT3/tSTAT3 protein ratios from blot (A) at 15 minutes and 3 hours post-stimulation. Measurements are in arbitrary units, presented as mean ± standard deviation (SD). Statistical significance determined by one-way ANOVA with 5% FDR (Benjamini–Hochberg method) gave no significant comparisons. Data derived from N=3 neurotypical male hiPSC cell lines, each with one technical replicate. Data points are colour-coded per donor line: red for M3_CTR, blue for 127_CTM, and green for 014_CTM.

**Supplementary Table 3 -** Two-way ANOVA of quantified western blotting pSTAT3/tSTAT3 signal of treated NPC in mono-culture at D18 of differentiation from N=3 donors, with 100ng/ml IL-6 or acetic acid vehicle, plus different concentrations of recombinant IL-6Ra (1, 10, 25, 50 and 100ng/ml).

| Source of Variation | DF | F (DFn, DFd) | % of total variation | P value | P value summary |
|---|---|---|---|---|---|
| **Interaction** | 4 | F (4, 20) = 31.94 | 17.50 | <0.0001 | **** |
| **[rIL6Ra] Factor** | 4 | F (4, 20) = 36.55 | 20.03 | <0.0001 | **** |
| **Treatment Factor** | 1 | F (1, 20) = 436.0 | 59.73 | <0.0001 | **** |

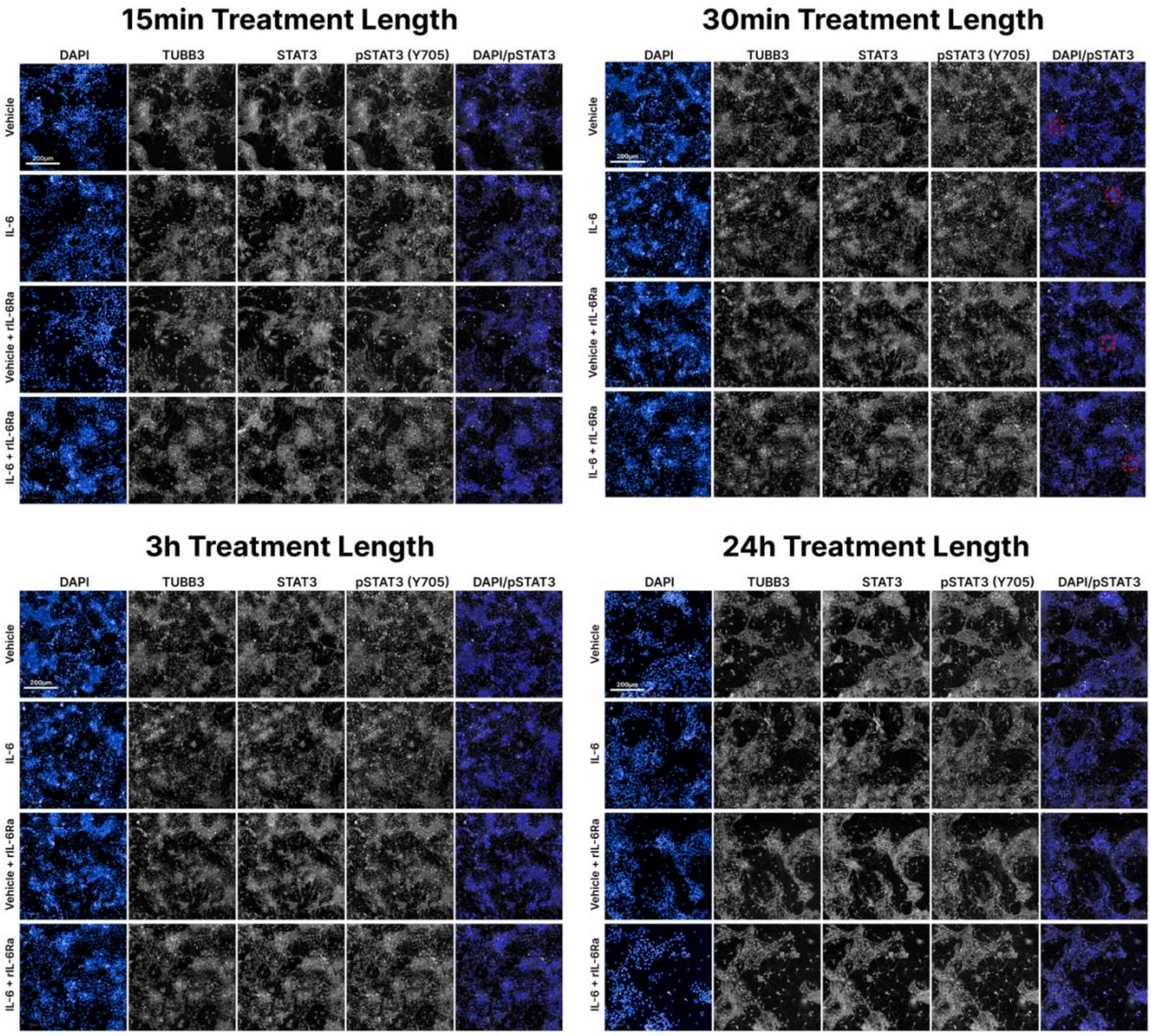


**Supplementary Figure 8 -** Image Panels for pSTAT3 and total STAT3 immunocytochemistry staining in D24 hiPSC-derived NPCs from donor line 127_CTM. Cells were treated with Vehicle (Acetic acid), IL-6 100ng/ml, Vehicle + rIL6Ra 100ng/ml or IL-6 + rIL6Ra, then stained for TUBB3 (#TUJ Aves Labs), STAT3 (Cell Signalling #9139) and phospho-STAT3 (Cell Signalling #9145) and imaged at 20x. Scale bar represents 200µm. Red dotted boxes in the 30min treatment panel correspond to the locations from which digital zoom box images were taken for Figure 1E.

**Supplementary Table 4 –** Number of NPCs measured per time and treatment condition of the immunocytochemistry experiment presented in Figure 1E, F and Supplementary Figure 7.

| Treatment | Time | Total Sum of Cells Measured |
|---|---|---|
| **IL-6** | 15min | 27263 |
| | 24h | 26827 |
| | 30min | 18525 |
| | 3h | 23192 |
| **IL-6 + rIL-6Ra** | 15min | 17733 |
| | 24h | 21633 |
| | 30min | 17008 |
| | 3h | 18413 |
| **Vehicle** | 15min | 27436 |
| | 24h | 25733 |
| | 30min | 22716 |
| | 3h | 22913 |
| **Vehicle + rIL-6Ra** | 15min | 21412 |
| | 24h | 23774 |
| | 30min | 19557 |
| | 3h | 21320 |

**Supplementary Table 5 – Sanger Sequencing of sIL6Ra Asp358Ala SNP.** All lines showed homogeneous protective C/C genotype.

| Line | M3_CTR | 014_CTM | 127_CTM | 069_CTF |
|---|---|---|---|---|
| **Sanger Sequencing Trace** | 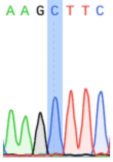  | 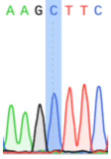  | 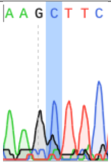  | 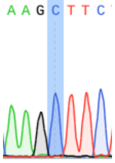  |
| **Asp358Ala (A > C) Genotype** | C/C | C/C | C/C | C/C |

**Supplementary Table 6 -** List of cytokine targets or controls at each coordinate dot on membranes of the Proteome Profiler Human XL Cytokine Array kit from R&D systems. Mean signal values from cytokine profiler dot blots, including all 105 cytokines, averaged from two replicate dots that were then backgrounded to negative reference dots and normalised to positive control reference dots. Table cells are coloured on gradient from low (white) to high (red). * IL-6 spiked in treatment groups. Grid coordinate corresponds to grid in Supplementary Figure 4.

| Cytokine | Array Coordinate | Control Vehicle | Control IL-6 |
|---|---|---|---|
| Adiponectin | **A3-4** | 0.108 | 0.131 |
| Apolipoprotein A-I | **A5-6** | 0.124 | 0.138 |
| Angiogenin | **A7-8** | 0.176 | 0.178 |
| Angiopoietin-1 | **A9-10** | 0.085 | 0.087 |
| Angiopoietin-2 | **A11-12** | 0.097 | 0.116 |
| BAFF | **A13-14** | 0.072 | 0.075 |
| BDNF | **A15-16** | 0.070 | 0.083 |
| Complement Component C5/C5a | **A17-18** | 0.055 | 0.072 |
| CD14 | **A19-20** | 0.125 | 0.156 |
| CD30 | **A21-22** | 0.065 | 0.068 |
| CD40 ligand | **B3-4** | 0.104 | 0.132 |
| Chitinase 3-like 1 | **B5-6** | 0.773 | 0.840 |
| Complement Factor D | **B7-8** | 0.116 | 0.108 |
| C-Reactive Protein | **B9-10** | 0.099 | 0.103 |
| Cripto-1 | **B11-12** | 0.079 | 0.078 |
| Cystatin C | **B13-14** | 0.118 | 0.172 |
| Dkk-1 | **B15-16** | 0.070 | 0.078 |
| DPPIV | **B17-18** | 0.093 | 0.122 |
| EGF | **B19-20** | 0.067 | 0.064 |
| EMMPRIN | **B21-22** | 0.156 | 0.161 |
| ENA-78 | **C3-4** | 1.135 | 1.414 |
| Endoglin | **C5-6** | 0.111 | 0.139 |
| Fas Ligand | **C7-8** | 0.113 | 0.099 |
| FGF basic | **C9-10** | 0.098 | 0.110 |
| FGF-7 | **C11-12** | 0.080 | 0.081 |
| FGF-19 | **C13-14** | 0.112 | 0.145 |
| Flt-3 Ligand | **C15-16** | 0.067 | 0.075 |
| G-CSF | **C17-18** | 0.071 | 0.067 |
| GDF-15 | **C19-20** | 0.106 | 0.102 |
| GM-CSF | **C21-22** | 0.589 | 0.574 |
| GROα | **D1-2** | 0.115 | 0.241 |

| | | | |
|---|---|---|---|
| Growth Hormone | **D3-4** | 0.093 | 0.121 |
| HGF | **D5-6** | 0.101 | 0.111 |
| ICAM-1 | **D7-8** | 0.130 | 0.118 |
| IFN-γ | **D9-10** | 0.097 | 0.094 |
| IGFBP-2 | **D11-12** | 0.546 | 0.596 |
| IGFBP-3 | **D13-14** | 0.064 | 0.075 |
| IL-1α | **D15-16** | 0.073 | 0.077 |
| IL-1β | **D17-18** | 0.070 | 0.069 |
| IL-1ra | **D19-20** | 0.120 | 0.107 |
| IL-2 | **D21-22** | 0.077 | 0.074 |
| IL-3 | **D23-24** | 0.063 | 0.061 |
| IL-4 | **E1-2** | 0.111 | 0.145 |
| IL-5 | **E3-4** | 0.098 | 0.114 |
| IL-6* | **E5-6** | 0.098 | 0.788 |
| IL-8 | **E7-8** | 1.178 | 1.269 |
| IL-10 | **E9-10** | 0.084 | 0.102 |
| IL-11 | **E11-12** | 0.090 | 0.108 |
| IL-12 p70 | **E13-14** | 0.072 | 0.071 |
| IL-13 | **E15-16** | 0.059 | 0.072 |
| IL-15 | **E17-18** | 0.066 | 0.073 |
| IL-16 | **E19-20** | 0.060 | 0.063 |
| IL-17A | **E21-22** | 0.118 | 0.133 |
| IL-18 | **E23-24** | 0.068 | 0.062 |
| IL-19 | **F1-2** | 0.108 | 0.123 |
| IL-22 | **F3-4** | 0.125 | 0.144 |
| IL-23 | **F5-6** | 0.103 | 0.109 |
| IL-24 | **F7-8** | 0.139 | 0.137 |
| IL-27 | **F9-10** | 0.091 | 0.104 |
| IL-31 | **F11-12** | 0.077 | 0.091 |
| IL-32 | **F13-14** | 0.066 | 0.083 |
| IL-33 | **F15-16** | 0.061 | 0.072 |
| IL-34 | **F17-18** | 0.419 | 0.381 |
| IP-10 | **F19-20** | 0.065 | 0.074 |
| I-TAC | **F21-22** | 0.063 | 0.071 |
| Kallikrein 3 | **F23-24** | 0.077 | 0.068 |
| Leptin | **G1-2** | 0.115 | 0.129 |

| | | | |
|---|---|---|---|
| LIF | **G3-4** | 0.112 | 0.091 |
| Lipocalin-2 | **G5-6** | 0.116 | 0.081 |
| MCP-1 | **G7-8** | 0.538 | 0.575 |
| MCP-3 | **G9-10** | 0.611 | 0.709 |
| M-CSF | **G11-12** | 0.085 | 0.111 |
| MIF | **G13-14** | 0.228 | 0.225 |
| MIG | **G15-16** | 0.060 | 0.084 |
| MIP-1α/MIP-1β | **G17-18** | 0.102 | 0.424 |
| MIP-3α | **G19-20** | 0.066 | 0.090 |
| MIP-3β | **G21-22** | 0.066 | 0.069 |
| MMP-9 | **G23-24** | 1.301 | 1.341 |
| Myeloperoxidase | **H1-2** | 0.138 | 0.142 |
| Osteopontin | **H3-4** | 1.173 | 1.112 |
| PDGF-AA | **H5-6** | 0.128 | 0.089 |
| PDGF-AB/BB | **H7-8** | 0.114 | 0.103 |
| Pentraxin 3 | **H9-10** | 0.444 | 0.477 |
| PF4 | **H11-12** | 0.064 | 0.102 |
| RAGE | **H13-14** | 0.067 | 0.092 |
| RANTES | **H15-16** | 0.061 | 0.129 |
| RBP-4 | **H17-18** | 0.057 | 0.082 |
| Relaxin-2 | **H19-20** | 0.062 | 0.080 |
| Resistin | **H21-22** | 0.072 | 0.107 |
| SDF-1α | **H23-24** | 0.072 | 0.099 |
| Serpin E1 | **I1-2** | 0.295 | 0.297 |
| SHBG | **I3-4** | 0.139 | 0.160 |
| ST2 | **I5-6** | 0.101 | 0.098 |
| TARC | **I7-8** | 0.107 | 0.117 |
| TFF3 | **I9-10** | 0.081 | 0.104 |
| TfR | **I11-12** | 0.084 | 0.100 |
| TGF-α | **I13-14** | 0.065 | 0.089 |
| Thrombospondin-1 | **I15-16** | 0.059 | 0.076 |
| TNF-α | **I17-18** | 0.062 | 0.105 |
| uPAR | **I19-20** | 0.205 | 0.292 |
| VEGF | **I21-22** | 0.074 | 0.093 |
| VCAM-1 | **J11-12** | 0.080 | 0.125 |
| Vitamin D | **J5-6** | 0.129 | 0.114 |

| | | | |
|---|---|---|---|
| CD31 | **J7-8** | 0.143 | 0.144 |
| TIM-3 | **J9-10** | 0.100 | 0.127 |

**Supplementary Table 7 -** Two-way ANOVA of MSD quantified IL-8, MIP-1α, TNF α and VEGF secretion from D14 MGLs and D18 NPCs from N=3 donors in monoculture treated for 24h with 100ng/ml IL-6.

| Cytokine | Source of Variation | DF | F (DFn, DFd) | P Value | P Value Summary |
|---|---|---|---|---|---|
| **IL-8** | Interaction | 1 | F (1, 8) = 1.119 | 0.3211 | ns |
| | Treatment Factor | 1 | F (1, 8) = 1.140 | 0.3169 | ns |
| | Cell Factor | 1 | F (1, 8) = 4845 | <0.0001 | **** |
| **MIP-1α** | Interaction | 1 | F (1, 8) = 0.1479 | 0.7106 | ns |
| | Treatment Factor | 1 | F (1, 8) = 0.1479 | 0.7106 | ns |
| | Cell Factor | 1 | F (1, 8) = 11.55 | 0.0094 | ** |
| **VEGF** | Interaction | 1 | F (1, 8) = 0.1200 | 0.7379 | ns |
| | Treatment Factor | 1 | F (1, 8) = 0.1258 | 0.7320 | ns |
| | Cell Factor | 1 | F (1, 8) = 33.52 | 0.0004 | *** |
| **TNF-α** | Interaction | 1 | F (1, 8) = 6.255 | 0.0369 | * |
| | Treatment Factor | 1 | F (1, 8) = 6.212 | 0.0374 | * |
| | Cell Factor | 1 | F (1, 8) = 49.03 | 0.0001 | *** |

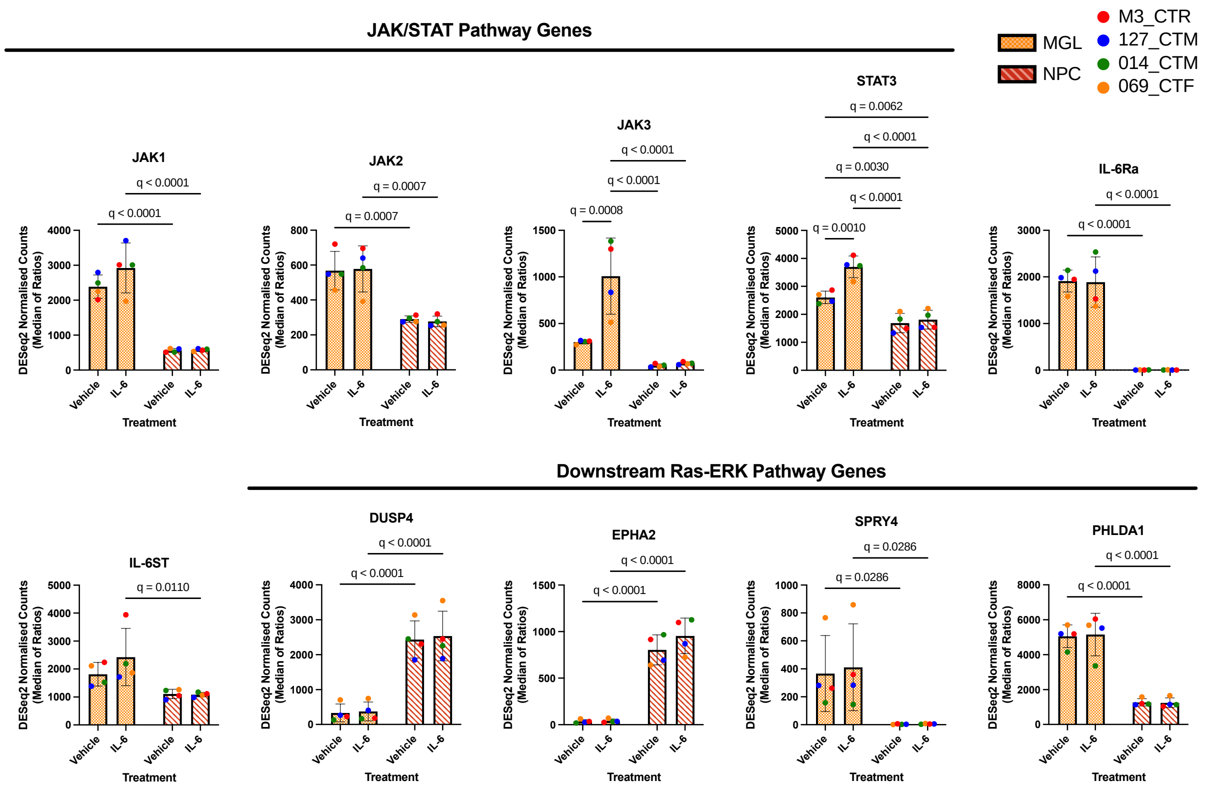


**Supplementary Figure 9 –** Counts of genes involved in IL-6 signalling from both MGL and NPC RNAseq datasets, as well as genes downstream of the Ras-ERK pathway (Chesnokov, Yadav and Chefetz, 2022). Units are normalised by the DESeq2 package as a median of ratios. Significant adjusted p-values after FDR correction using the BH method are labelled on the graphs.

**Supplementary Table 8 –** List of common genes in both up-regulated IL-6 MGL response and those up-regulated in post-mortem tissue from SZ patients.

| Gene Sets | Common Genes in Both Sets |
|---|---|
| Signature A and SZ Upregulated Genes | BATF, CA12, CHI3L2, CLEC4E, FPR2, GK, GRAMD1A, JAK3, KREMEN1, MCOLN2, MT2A, MYO1G, S100A8, SOCS3, SOD2, STAT3, ZC3H12A |

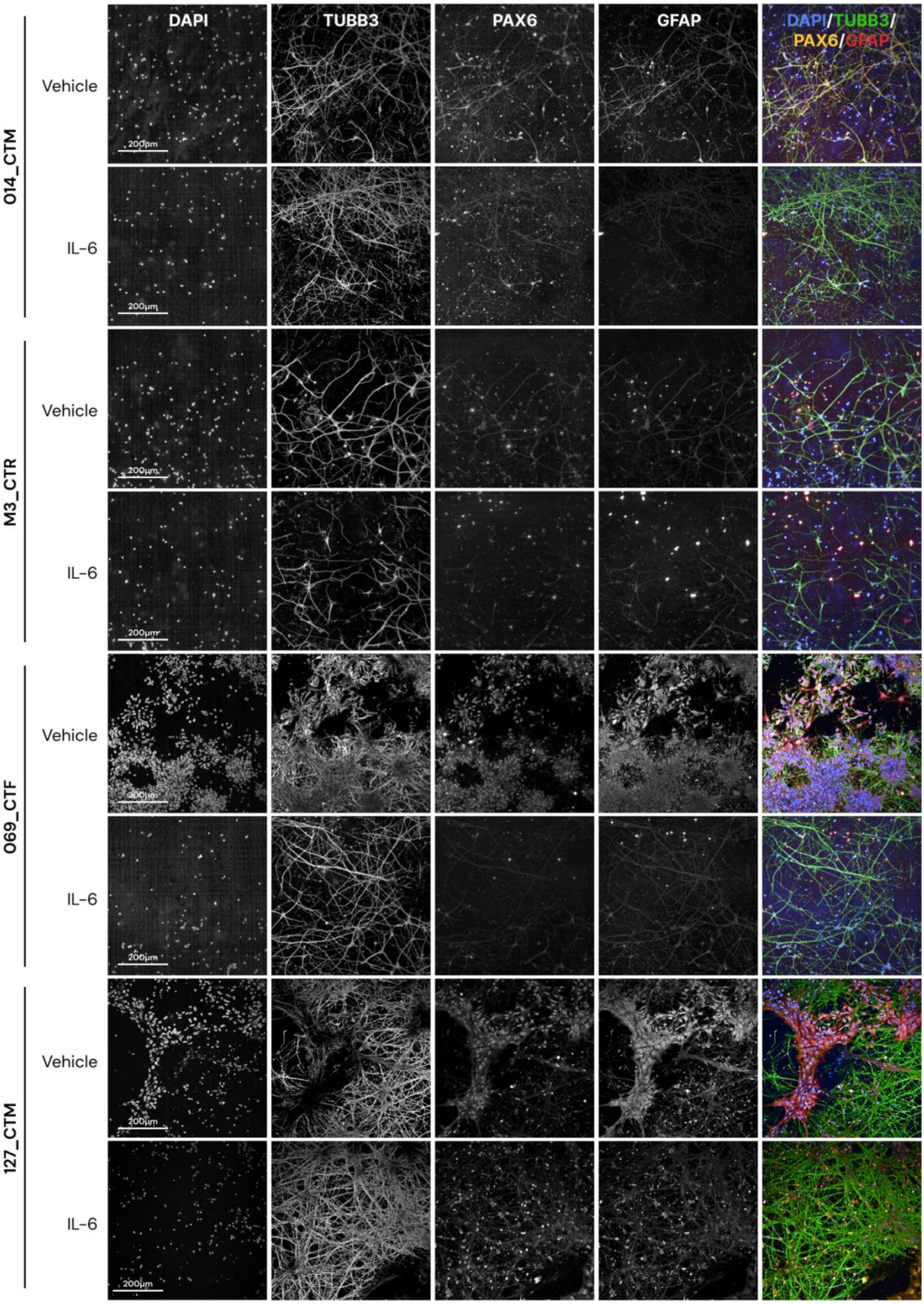


**Supplementary Figure 10 – Representative images of neuronal marker expression for all four donor lines.** Variation in cell type differentiation is evident across each donor line. These are represented in the percentages of cell type combinations found in the following supplementary table 6. Scale bar represents 200µm.

**Supplementary Table 9 -** Cell populations as a percentage of total number of cells counted within each donor (N = 4 donors averaged from N= 4 technical well culture repeats each), averaged across all images to one single percentage per donor. These percentages correspond to those plotted in figure 5B.

| Cell Type | Donor | Vehicle Condition (cell %) | IL-6 Condition (cell %) |
|---|---|---|---|
| **TUBB3+/PAX6-/GFAP-** | M3_CTR | 81.82 | 84.59 |
| | 014_CTM | 86.89 | 84.72 |
| | 069_CTF | 80.87 | 90.98 |
| | 127_CTM | 40.98 | 60.62 |
| **TUBB3+/PAX6+/GFAP-** | M3_CTR | 0 | 0.05 |
| | 014_CTM | 0.04 | 0 |
| | 069_CTF | 0.05 | 0 |
| | 127_CTM | 16.82 | 10.95 |
| **TUBB3+/Pax6-/GFAP+** | M3_CTR | 11.81 | 11.57 |
| | 014_CTM | 11.68 | 14.52 |
| | 069_CTF | 13.01 | 5.67 |
| | 127_CTM | 38.97 | 26.49 |
| **TUBB3+/Pax6+/GFAP+** | M3_CTR | 0.07 | 0 |
| | 014_CTM | 0 | 0.02 |
| | 069_CTF | 0 | 0 |
| | 127_CTM | 0.07 | 0 |
| **TUBB3-/Pax6+/GFAP+** | M3_CTR | 0.15 | 0 |
| | 014_CTM | 0 | 0 |
| | 069_CTF | 0 | 0 |
| | 127_CTM | 0.01 | 0.33 |
| **TUBB3-/Pax6+/GFAP-** | M3_CTR | 5.62 | 3.29 |
| | 014_CTM | 1.3 | 0.72 |
| | 069_CTF | 6 | 3.35 |
| | 127_CTM | 1.18 | 1.05 |
| **TUBB3-/Pax6-/GFAP+** | M3_CTR | 0.53 | 0.5 |
| | 014_CTM | 0.09 | 0.03 |
| | 069_CTF | 0.08 | 0 |
| | 127_CTM | 1.97 | 0.59 |

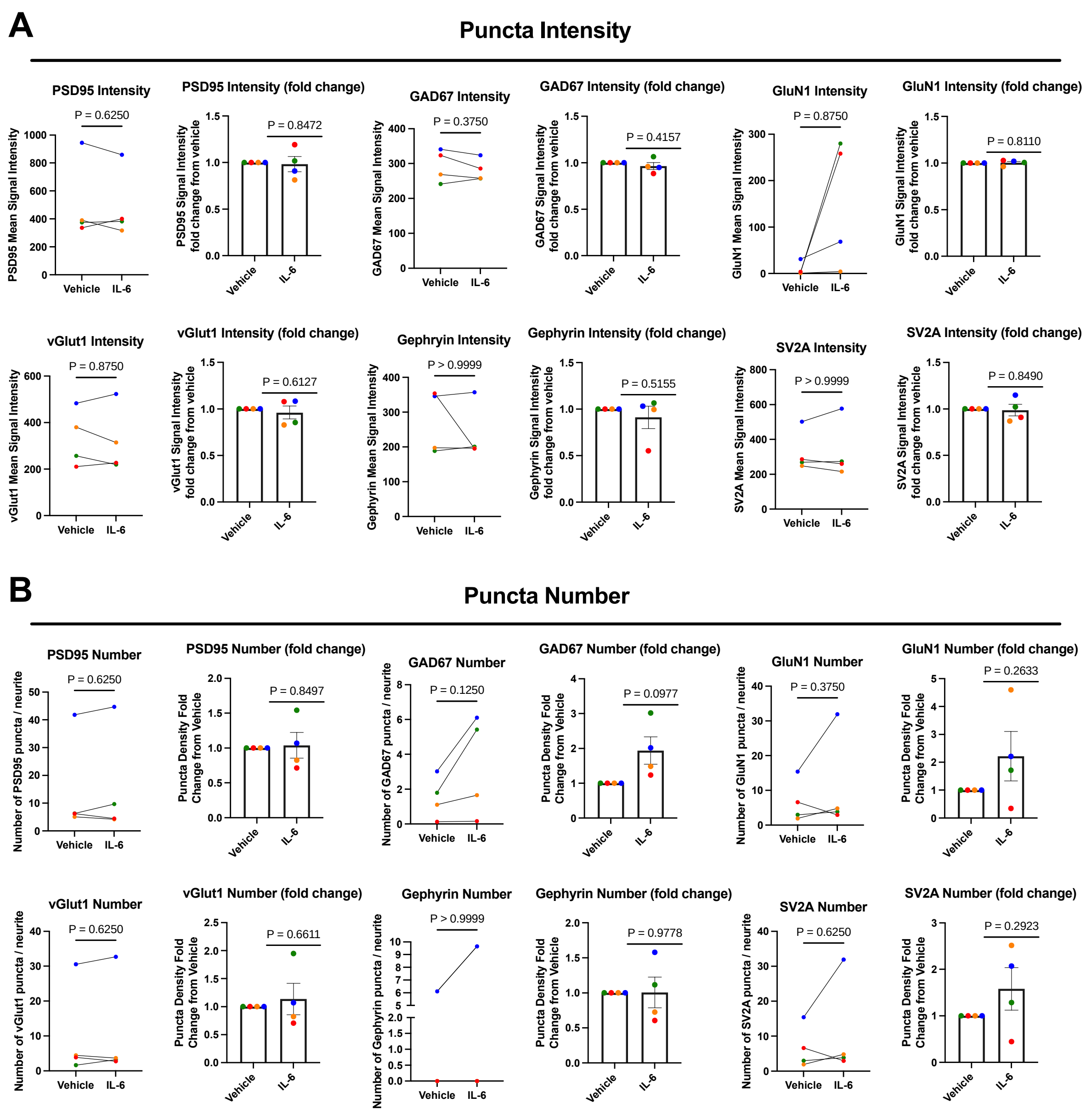


**Supplementary Figure 11 – Raw and fold change comparisons of puncta assay.** Puncta were identified and measured for stain intensity (A) and number per neurite (identified by MAP2 staining) (B). Raw values are plotted as line graphs and fold changes as bar graphs. Given the non-parametric nature of the data, a Wilcoxon matched-pairs signed rank tests was used to compare raw values from vehicle to treated samples. After fold change calculation to the vehicle, a one-sided Wilcoxon test was used to compare the fold change from vehicle of intensity and puncta number. The p-value for each of these tests is formatted on each graph. Bar graphs plotted as mean with standard deviation (SD) error bars, and points coloured by donor line: red (M3_CTR), blue (127_CTM), green (014_CTM) and orange (069_CTF), using a total of N=4 control donors as biological replicates.
